# An extra-glycolytic function for hexokinase 2 as an RNA-binding protein regulating *SOX10* mRNA translation in melanoma

**DOI:** 10.1101/2024.08.13.607712

**Authors:** Ana Luisa Dian, Antoine Moya-Plana, Giuseppina Claps, Céline M. Labbé, Virginie Quidville, Dorothée Baille, Séverine Roy, Virginie Raynal, Sylvain Baulande, Caroline Robert, Stéphan Vagner

## Abstract

Several studies have reported the importance of aerobic glycolysis in melanoma development. Although metabolic benefits of glycolysis have been extensively described in tumor cells, the extra-metabolic functions linked to this energetic pathway in melanoma growth and proliferation have not been clearly established yet. Recently, some key glycolytic enzymes, such as GAPDH and PKM2, were reported to regulate mRNA translation. Translational control of gene expression is considered as a critical effector in cancer biology, representing a highly promising area of research. Here, we report that Hexokinase 2 (HK2), a glucose kinase that catalyzes the first step of glycolysis, is an RNA binding protein (RBP) that regulates mRNA translation in melanoma. We show that siRNA-mediated HK2 depletion changes the translational landscape of melanoma cells. Polysome profiling experiments and RNA-Seq indicate that the translational regulation exerted by HK2 is partly independent of the metabolic status or the glycolytic pathway. We found that HK2 specifically regulates the translation of the mRNA encoding SOX10, a transcription factor implicated in the regulation of tumor initiation, maintenance and progression in melanoma. RNA-protein interaction assays, including crosslinking immunoprecipitation (CLIP), indicate that HK2 is an RBP whose interaction with RNA is independent of its hexokinase activity or subcellular localization. We also show that HK2 specifically associates with the 5’ untranslated region (5’UTR) of the *SOX10* mRNA, and that several deletions in this region decreases both HK2-*SOX10* mRNA association and *SOX10* 5’ UTR-mediated translation. We further show that HK2-dependent SOX10 translational regulation is involved in melanoma cell proliferation and colony formation. Collectively, our data highlight a non-metabolic function of HK2, indicating that melanoma cells may enhance glycolysis for purposes beyond simple anabolism.

## INTRODUCTION

Aerobic glycolysis, in which glucose is converted into lactate, is a key metabolic pathway in tumorigenesis. This pathway is usually promoted by cancer cells even when local conditions are suitable to utilize the oxidative phosphorylation (OXPHOS), a more efficient mechanism to generate ATP. This specific metabolic process is called the Warburg effect (Warburg, 1956; DeBerardinis & Chandel, 2020). Although metabolic benefits of glycolysis have been established for sustaining biosynthesis and cellular proliferation in tumor cells (Lunt & Vander Heiden, 2011; Vander Heiden *et al*, 2009) some data suggest that extra-metabolic functions are linked to this energetic pathway.

Over the past decades, several enzymes involved in the intermediary metabolism were shown to directly interact with RNAs (Castello *et al*, 2015). Indeed, several metabolic enzymes were found to bind RNA in the vicinity of their substrate-binding pockets (Castello *et al*, 2016; Nagy & Rigby, 1995). One example is the Rossmann fold, a di-nucleotide domain typically associated with the binding of nucleotides such as the nicotinamide adenine dinucleotide (NAD) (Rossman *et al*, 1975). This fold plays a crucial role in the function of many enzymes, particularly dehydrogenases such as GAPDH, which catalyze oxidation-reduction reactions. Interestingly, this same domain has been implicated in mediating RNA-binding (Nagy & Rigby, 1995).

Beyond dehydrogenases, several other enzymatic activities are found in the glycolytic pathway including kinases for glucose (HK1/2), pyruvates (PKM1/2), and enolases (ENO1/2/3). While PKM2 and ENO1 have been found to bind RNAs, the possible RNA-binding activity of hexokinase 2 (HK2), the first and limiting glycolysis enzyme catalyzing the irreversible phosphorylation of glucose to form glucose-6-phosphate, has never been addressed. High levels of HK2 expression have been reported in many different types of cancer, and this upregulation is typically associated with poor outcomes in patients (Wang *et al*, 2017; Wolf *et al*, 2011; Patra *et al*, 2013; Kudryavtseva *et al*, 2016; Liu *et al*, 2013). Although the role of HK2 in tumorigenesis has been attributed to its glycolytic activity, HK2 has also been shown to execute non-canonical functions that often regulate processes that are highly relevant for cell transformation and cancer development. In particular, Blaha *et al*. (2022) demonstrated a novel non-catalytic mechanism by which HK2 contributes to SNAIL-mediated EMT and metastasis in breast cancer (Blaha *et al*, 2022). Intriguingly, HK2 has also been identified as a candidate RBP by orthogonal large-scale quantitative methods (Trendel *et al*, 2019; Queiroz *et al*, 2019; Urdaneta *et al*, 2019; Backlund *et al*, 2020) and many glucose-binding proteins were recently reported to bind RNA in human cell lines (Miao *et al*, 2023). Together with recent evidences showing that some key-glycolytic enzymes, such as GAPDH (glyceraldehyde-3-phosphate dehydrogenase) and PKM (pyruvate kinase) were able to regulate the translation of mRNAs (Simsek *et al*, 2017; Chang *et al*, 2013) in human T-cells and mouse embryonic stem cells respectively, we explored the role of HK2 as an RBP regulating mRNA translation in human cancer cells.

Here we describe a novel extra-glycolytic function of HK2 in melanoma cell lines chosen due to the critical role of glycolysis and OXPHOS in melanoma development (Scott *et al*, 2011; Qin *et al*, 2010). We demonstrate that the HK2 glucose kinase is a *bona-fide* RBP that regulates the translation of the *SOX10* mRNA, thereby regulating cell proliferation.

## RESULTS

### HK2 regulates mRNA translation

Upon stress stimuli and/or acquisition of mutations, melanocytes evolve from early superficial melanoma with radial growth (RGP) to early invasive melanoma (VGP) and metastatic tumor. We evaluated the level of HK2 in a variety of melanoma cell lines. We observed increased levels of HK2 in invasive cell lines (A375, WM35, SKMel10) compared to VGP (WM793), RGP (SBCL2) or normal human melanocytes (Figure S1A). In addition, in patients with cutaneous melanoma, HK2 expression was significantly higher in stages III-IV compared to stages I-II (p=0.026) (Figure S1B).

Since PKM2 and GAPDH were shown to associate with polysomes (Kejiou *et al*, 2023; Simsek *et al*, 2017), we analyzed whether HK2 could also be associated with polysomes. To this end, we performed western blot analysis on polysome fractions isolated from A375 cells, one of the melanoma cell lines with the highest level of HK2 (Figure S1C). We found that, as for PKM2 and GAPDH, HK2 is present in ribosome-containing fractions (Figure 1A). This is specific since another glycolytic enzyme, LDHB, is not associated to ribosome-containing fractions.

**Figure 1.**
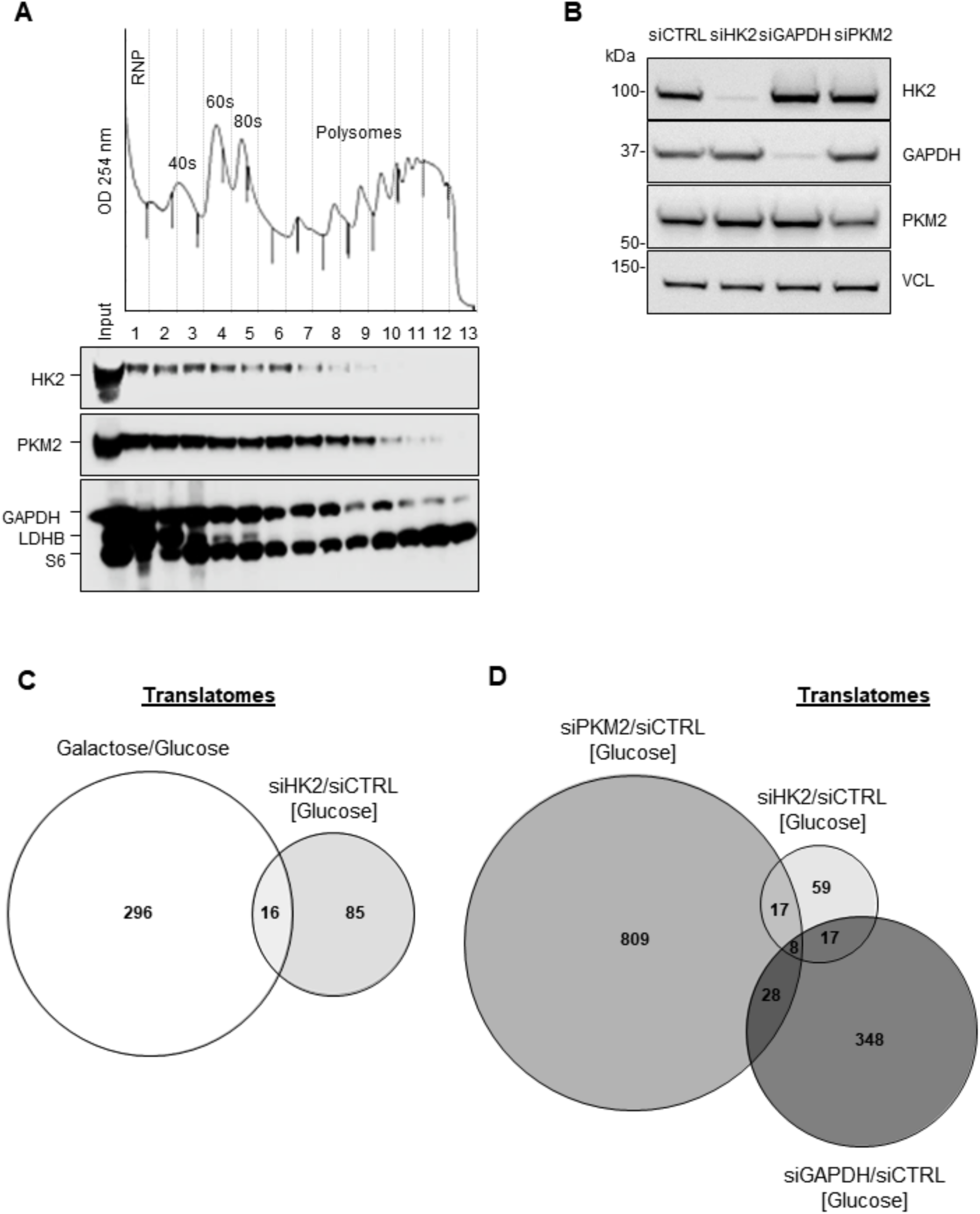
HK2 regulates mRNA translation in melanoma cells. **(A)** Western blot analysis of HK2, PKM2, GAPDH, LDHB and the ribosomal protein S6 in fractions (horizontal axes) obtained by sucrose-gradient (10-50%) ultracentrifugation of lysates from A375 cells. The cells were treated with 100 μg/ml Cycloheximide for 10 min prior to lysis to stabilize polysomes. Nucleic acids were monitored by OD254 nm, and proteins were monitored in each fraction by western blot. **(B)** Western blot analysis of HK2, GAPDH and PKM2 protein levels in A375 melanoma cells upon siRNA-mediated depletion of the enzymes (siHK2, siGAPDH and siPKM2, respectively) or control (siCTRL). VCL was used as loading control. **(C)** Comparison of translatomes between A375 cells cultured in galactose-containing medium relative to glucose (312 deregulated mRNAs, 214 being upregulated and 98 downregulated) and siRNA-mediated depletion of HK2 in cells cultured in glucose-containing medium relative to control (siHK2/siCTRL; 101 deregulated mRNAs, being 52 upregulated and 49 downregulated) (n = 3 biological replicates). **(D)** Comparison of translatomes between A375 cells upon siRNA-mediated depletion of key glycolytic enzymes (HK2, GAPDH, PKM2) in A375 cells relative to control. 401 mRNA candidates (269 upregulated and 132 downregulated) were found deregulated upon siRNA-mediated depletion of GAPDH relative to control (siGAPDH/siCTRL), and 862 mRNA candidates (554 upregulated and 308 downregulated) were found deregulated upon siRNA-mediated depletion of PKM2 (siPKM2/siCTRL). P-value ≤ 0.05 with fold change > 0.7 (DESeq2).

To investigate whether HK2 is involved in the translational regulation of specific mRNAs, we depleted (siRNA) HK2 in A375 melanoma cells. Of note, depletion of HK2 did not lead to a decrease in the levels of GAPDH and PKM2 (Figure 1B). We used RNA sequencing to simultaneously quantify the abundance of total transcripts and those that are being actively translated, i.e. that are associated with polysomes (>4 ribosomes). The analysis of mRNAs that are differentially recruited to heavy polysome fractions in HK2-depleted cells under normoglucidic conditions showed that the translation status of 101 mRNAs was changed (log2(fold change) > 0.7; pvalue < 0.05) (Figure 1C and Table S1).

To determine if this translational regulation is linked to the metabolic status, we cultured A375 cells in medium containing galactose instead of glucose for 24 h. The metabolism of galactose eventually converges with the glucose metabolism through the Leloir pathway. However, because this conversion occurs at a much slower rate than glucose entry into glycolysis, culture with galactose favours oxidative phosphorylation (OXPHOS) (Zhang *et al*, 2013; Wang *et al*, 2018; Weinberg *et al*, 2010). To confirm that A375 cells switch their metabolic profile in the presence of galactose, we employed the Seahorse metabolic flux assay to measure the oxygen consumption rate (OCR) and the extra-cellular acidification rate (ECAR) of living cells in culture. We observed that the ECAR, an indicator of aerobic glycolysis, was significantly higher in A375 cells cultured in glucose-containing medium compared to cells cultured in galactose. On the other hand, the OCR, an indicator of OXPHOS, was higher than ECAR in cells cultured in galactose-containing medium (Figure S2). These results therefore confirmed the metabolic switch of the melanoma cells from glycolysis to mitochondrial respiration. We found that modifying the metabolic status of cells by growing them in the presence of galactose instead of glucose impacted the translation of 312 mRNAs (Figure 1C and Table S1). Only about 20% of mRNAs regulated by HK2 are also regulated by galactose indicating that translation regulation by HK2 is partly independent of the metabolic status of the cells (Figure 1C). We also compared the subsets of mRNAs translationally regulated by HK2, PKM2 and GAPDH. We found that 59 mRNAs are specifically regulated by HK2, but not by the other tested glycolytic enzymes (i.e. PKM2 or GAPDH) (Figure 1D).

In parallel, we performed RT-qPCR (reverse transcription quantitative Polymerase Chain Reaction) on a panel of 84 cancer-relevant mRNAs (Table S2). By comparing the presence of mRNAs on heavy polysomes fractions according to HK2 expression (siHK2 vs siCTRL), we identified 6 mRNAs that were specifically translationally downregulated upon siRNA-mediated HK2 depletion (Figure S3A-B). Among them, we were particularly interested on the *SOX10* mRNA that encodes the Sex Determining Region Y (SRY)-Box Transcription Factor 10 (SOX10) because of its well-known activity as a transcription factor regulating tumor initiation, maintenance and progression in melanoma (Han *et al*, 2018; Graf *et al*, 2014; Cronin *et al*, 2018). Of note, the *SOX10* mRNA was not found in the list of 101 candidate mRNAs regulated by HK2 (Table S1) because of lack of statistical significance (p=0.3). We compared SOX10 expression in different types of malignancies and normal tissues in human in the TCGA database and found that cutaneous melanoma highly expressed SOX10 compared to other cancers (Figure S4A). Moreover, we confirmed that the prognosis of melanoma was correlated to SOX10 expression. In a cohort of melanoma patients, the quartile low-SOX10 group had a significantly higher overall survival than the quartile high-SOX10 group (p=0.012) (Figure S4B). Interestingly, when the patients were stratified according to their mutational profile, SOX10 expression appeared to be a powerful prognostic factor of oncologic outcomes in patients with BRAF-mutated melanoma (as A375 cell line) while no prognostic value was observed for RAS, NF1 and triple-negative melanomas (Figure S4C). This could be explained by the fact that aerobic glycolysis is upregulated in human BRAF^V600^ melanoma cells via a transcriptional regulation of HK2 and GLUT1 (Parmenter *et al*, 2014). Thus, the transcriptional activation of HK2 by the MAPK/ERK pathway may lead to a translational activation of *SOX10*.

To confirm the *SOX10* mRNA translational regulation mediated by HK2, we monitored the level of *SOX10* mRNA isolated from all polysome fractions of A375 cells upon siRNA-mediated depletion of HK2. We found that *SOX10* mRNA shifted from heavy polysome fractions to lighter polysome fractions upon HK2 depletion, indicating that the depletion of the enzyme decreases the translation of the *SOX10* mRNA in melanoma cells (Figure 2A). Conversely, the translation of the *HPRT* mRNA (negative control) was not affected by HK2 depletion (Figure 2A). In agreement with our polysome data suggesting that HK2 is required for the translation of the *SOX10* mRNA, HK2 depletion decreased SOX10 protein levels, as observed in western blot analyses of different BRAF-mutated (V600E) melanoma cell lines (Figure 2B). This regulation appears to be specific to HK2 as the depletion of GAPDH did not affect SOX10 expression (Figure 2B). In addition, the *SOX10* mRNA level, which was evaluated by RT-qPCR, remained unchanged upon HK2 depletion (Figure 2C). Collectively, our results demonstrate that HK2 regulates the translation of cancer-relevant mRNAs, and more specifically of the *SOX10* mRNA.

**Figure 2.**
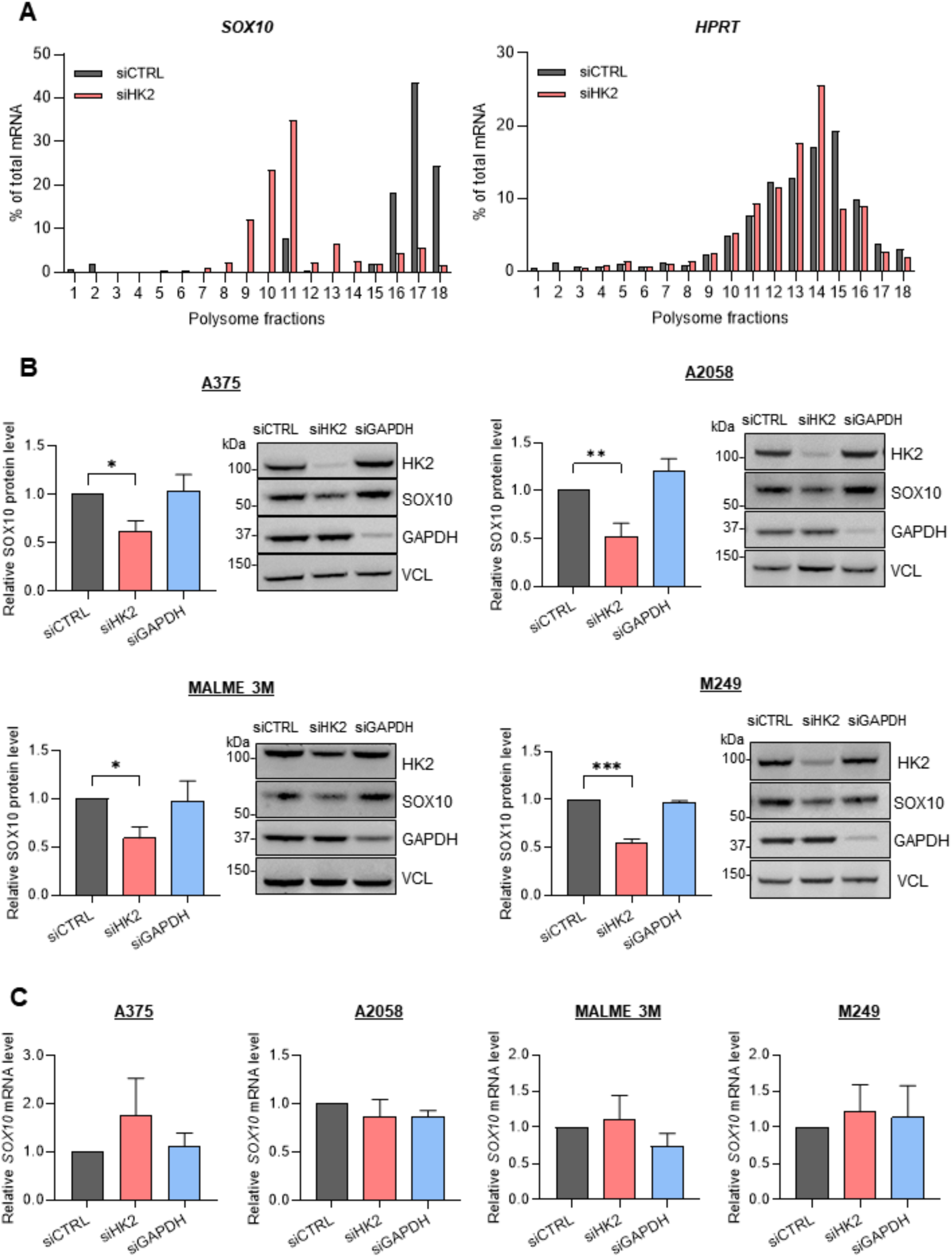
HK2 regulates the translation of the *SOX10* mRNA. **(A)** RT-qPCR quantification representing the % of *SOX10* mRNA (left panel) or *HPRT* mRNA (right panel) in fractions (horizontal axes) obtained by sucrose-gradient (10-50%) ultracentrifugation of lysates from A375 cells transfected with siRNAs targeting HK2 (siHK2, light red) or control (siCTRL, grey) (n = 2 biological replicates). **(B)** Western blot analysis of the SOX10 protein level in distinct melanoma cell lines transfected with siRNAs targeting HK2 (siHK2, light red), GAPDH (siGAPDH, light blue) or control (siCTRL, grey). The SOX10 protein quantification is normalized to VCL expression. *p*-values were calculated by two-tailed unpaired *t*-test (SD, n = 3 biological replicates) and only significant comparisons are shown (* p ≤ 0.05, ** p ≤ 0.01, *** p ≤ 0.001). **(C)** RT-qPCR quantification of the *SOX10* mRNA in different melanoma cell lines transfected with siRNAs targeting HK2 (siHK2, light red), GAPDH (siGAPDH, light blue) or control (siCTRL, grey). p-values were calculated by ordinary one-way ANOVA (SD, n = 3 biological replicates).

### HK2 directly binds RNA

To test the hypothesis that HK2 directly interacts with RNA, we first performed Complex Capture experiments (2C) (Asencio *et al*, 2018). This approach takes advantage of the ability of negatively charged nucleic acids to bind silica matrix-based columns, consequently retaining UV-crosslinked RNA-RBP complexes. HK2 was retained on the column in extracts of UV-C irradiated cells, but not detected in RNase A-treated samples, demonstrating that HK2 was retained on the column due to its UVC-induced covalent interaction with RNAs (Figure 3A). The well-characterized RBP HuR was used as a positive control in this assay, as well as the non-canonical RNA-metabolic enzymes GAPDH and PKM2. The metabolic enzymes ACOT1/2 and HSC70 were used as negative controls. To gain more evidence that HK2 binds to RNA, we performed CrossLinking ImmunoPrecipitation (CLIP) assay (Hafner *et al*, 2021; Lee & Ule, 2018). This assay relies on the UVC-dependent crosslinking of RNA-RBP complexes in living cells followed by cell lysis and partial RNA digestion with RNase I. The RNA molecules crosslinked to HK2 were then recovered through HK2 immunoprecipitation and radioactively labelled with ^32^P-yATP and T4 polynucleotide kinase (PNK). We observed a radioactive signal at the molecular size of HK2 in UVC-treated samples (Figure 3B). The signal was not detected in the non-crosslinked conditions or in IgG-immunoprecipitated conditions (negative controls) (Figure 3B). The signal was modulated according to the RNase I concentration used in UV-crosslinked cells; at higher concentrations, the radioactive signal collapsed to a sharp band at the molecular mass of HK2. These results indicate that HK2 directly interacts with RNA in living A375 melanoma cells.

**Figure 3.**
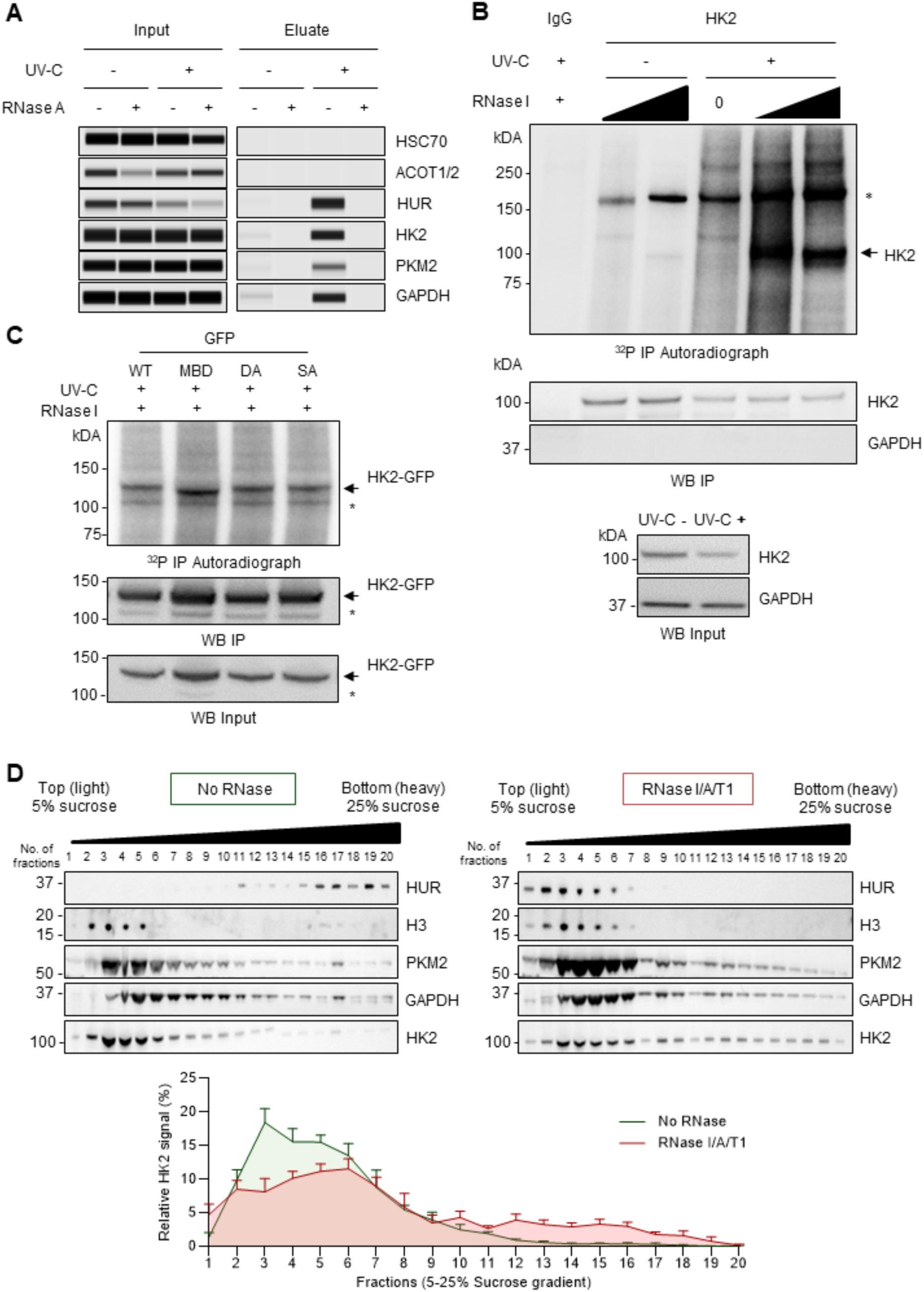
HK2 binds directly to RNA. **(A)** RNA-dependent retention of HK2 in silica matrix columns assessed by automated capillary western blot (Protein Simple ®) (n = 3 biological replicates). Specific HK2 signal was detected around the expected molecular mass in lysates of UV-C irradiated A375 cells not treated with RNase A prior to loading to the silica column. HUR, GAPDH and PKM2 were used as positive controls, and HSC70 and ACOT1/2 as negative controls. **(B)** CLIP from endogenous HK2 in A375 cells (n = 3 biological replicates). Upper panel: Autoradiography of HK2-RNA complexes in UV-C treated (+) or untreated (-) A375 cells. One major band is observed in the expected molecular mass of HK2, as indicated by a black arrow, and the asterisk (*) indicates a non-specific band. Middle and lower panel: western blots of HK2 and IgG immunoprecipitation, and HK2 and GAPDH (normalization control) input, respectively. Same conditions as the upper panel. **(C)** CLIP from GFP-HK2 transfected HEK293T cells (n=3 biological replicates). Upper panel: autoradiography of HK2-GFP-RNA complexes in UV-C treated (+) cells. Middle and lower panel: western blots of GFP and GFP immunoprecipitation, respectively. Same conditions as the upper panel. The arrow indicates the expected HK2-GFP molecular mass, and the asterisk (*) indicates a non-specific band. FL: full length; MBD: mitochondrial-binding deficient; DA: non-glucose-binding mutant; SA: catalytically inactive mutant. **(D)** Sucrose density gradient (5-25%) centrifugation and fractionation of A375 cell lysates treated with RNase I/A/T1 or left untreated (SD, n = 5 biological replicates). Upper panel: western blot of the sucrose-gradient fractions (horizontal axes) obtained after ultracentrifugation of lysates. HUR was used as a positive control, and H3 as a negative control. Lower panel: western blot quantification representing the % of HK2 in each sucrose fraction.

We next examined whether the RNA-binding activity of HK2 was dependent on its hexokinase activity or subcellular localization. We ectopically expressed GFP-tagged HK2, either wild-type (HK2-WT) or mutants in HEK293T cells and performed the CLIP assay. We used the HK2-MBD mutant, which carries a deletion of the first 15 amino acids that are critical for the association of HK2 with the outer membrane of the mitochondria through its interaction with the voltage-dependent anion-selective channel 1 (VDAC-1) (Wolf *et al*, 2016). This binding has been shown to enhance glycolysis due to privilege access of HK2 to newly synthesized ATP generated by the mitochondria. However, HK2 has been shown to alternate between cytoplasmic and mitochondrial-bound states in response to environmental and metabolic stress (Guo *et al*, 2022). The second tested mutant was the HK2-DA mutant, in which alanine is substituted to two aspartic acid residues in both the amino- and carboxy-terminal domains of HK2 (D209A/D657A). Mutations in both sites inhibit HK2 binding to glucose (Nawaz *et al*, 2018). Additionally, we tested whether the catalytic activity of HK2 is necessary for its binding to RNA. To this end, alanine was substituted for two serine residues in both the amino- and carboxy-terminal domains of HK2 (HK2-SA mutant; S155A/S603A) (Guo *et al*, 2022). As expected, we observed a sharp radioactive signal at the molecular size of HK2-FL-GFP (∼129 kDa), which corresponds to the upper band (Figure 3C). The lower band, labelled with an asterisk, corresponds to a non-specific signal. Strikingly, all mutants retained their ability to bind RNA, indicating that the interaction of HK2 with RNA is independent of its enzymatic activity, its ability to bind to glucose or its subcellular localization.

To further study HK2-RNA interaction, we subjected RNase-treated or untreated lysates of A375 cells to sucrose density gradient (5-25%) ultracentrifugation and fractionation. By these means, we found that 18% of the total cellular HK2 changed its distribution in the sucrose gradient following RNase treatment, indicating that the composition of HK2-containing complexes is partially dependent on RNA (Figure 3D). Intriguingly, complexes containing either GAPDH or PKM2 showed less sensitivity to RNase treatment (Figure S5). In contrast to HUR, HK2 presented an RNA-dependent shift to more dense fractions suggesting that the presence of RNA might prevent the formation of heavy HK2-containing complexes.

### HK2 associates with the *SOX10* mRNA

We next determined whether HK2 could specifically associate with the *SOX10* mRNA, which was identified to be translationally regulated by HK2. By performing HK2 immunoprecipitation (IP) followed by RT-qPCR (RNA ImmunoPrecipitation-RIP analysis), we identified an enrichment of the *SOX10* mRNA in HK2 IP over the control IP using IgG antibodies of the same isotype (Figure 4A). mRNAs of similar or higher abundance such as *GAPDH*, *TBP* and *actin* were less enriched in the HK2 IP indicating that HK2 specifically binds to the *SOX10* mRNA (Figure 4A and S7A). To delineate the HK2 binding site on the *SOX10* mRNA, *CatRAPID* fragment prediction (Armaos *et al*, 2021) was employed and revealed a significant binding site of HK2 to the *SOX10* sequence located between nucleotides 206 and 321. This region corresponds to the 5′ untranslated region (5′ UTR) of the *SOX10* mRNA (Figure 4B).

**Figure 4.**
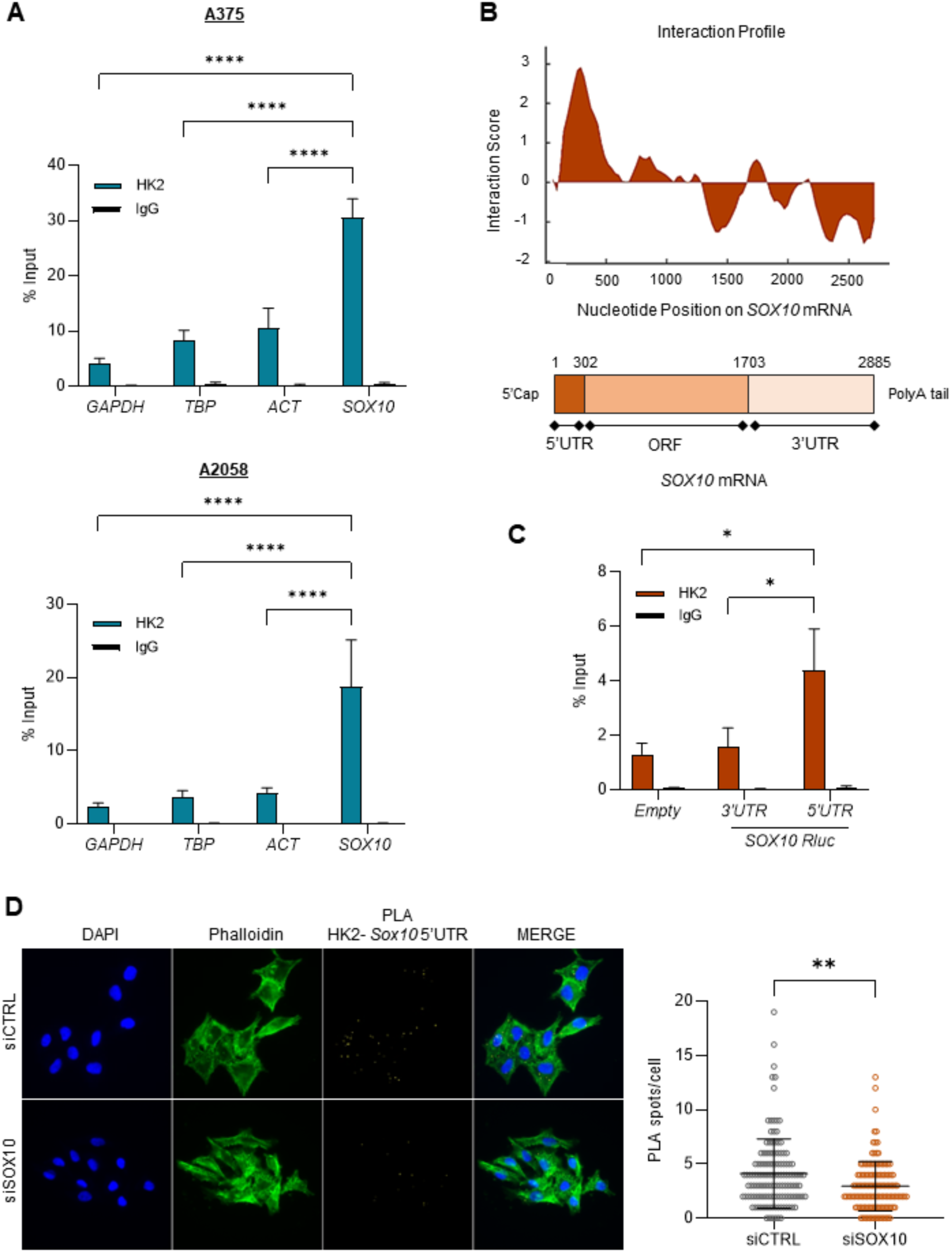
HK2 associates with the 5’UTR of the *SOX10* mRNA. **(A)** RIP experiment performed on A375 and A2058 melanoma cell lines (SD, n = 3 biological replicates). *SOX10* mRNA was analysed in IgG and HK2 immunoprecipitated samples and expressed as percentage of the mRNA present in the input. *GAPDH*, *TBP* and *ACT* mRNAs were used as negative controls. *p*-values were calculated by ordinary two-way ANOVA with Dunnett’s multiple comparisons test (SD, n = 3 biological replicates) (**** p ≤ 0.0001). **(B)** *In silico* prediction of HK2-*SOX10* mRNA interaction propensities using the *CatRAPID* algorithm. The highest interaction score observed between HK2 and the *SOX10* mRNA is at the *SOX10* 5’UTR. Upper panel: HK2-*SOX10* mRNA interaction profile. Lower panel: schematic representation of the *SOX10* mRNA. **(C)** RIP experiment performed on A375 cells transfected with luciferase reporters containing the *SOX10* 5’UTR and 3’UTR sequences upstream of the *Renilla* reporter gene. An Empty reporter was used as control. *Renilla* mRNA was analyzed in IgG and HK2 immunoprecipitated samples and expressed as percentage of the mRNA present in the input. *p*-values were calculated by ordinary two-way ANOVA with Dunnett’s multiple comparisons test (SEM, n = 6 biological replicates) (* p ≤ 0.05). **(D)** Representative fluorescence images and quantification of RNA-proximity ligation assays (PLA) of HK2 protein and *SOX10* mRNA in A375 cells transfected with siRNAs targeting SOX10 (siSOX10, orange) or control (siCTRL, grey) (n = 3 biological replicates). After 48h, cells were fixed, permeabilized, and incubated with anti-sense *SOX10* 5’UTR oligonucleotide probes and anti-HK2 antibody. Scale bar, 10 μm. HK2-*SOX10* mRNA PLA signal (yellow), Phalloidin staining (green), and DAPI staining (blue). The significance for PLA values was derived from the Mann-Whitney statistical test (SD, ** p ≤ 0.01).

We transfected A375 cells with plasmids containing either the *SOX10* 5’UTR, the 3’UTR or no *SOX10* sequences (empty vector) and performed RIP experiments. Consistent with the *in silico* prediction, we observed a significant enrichment of the 5’UTR-containing mRNA over the 3’UTR-containing mRNA (Figure 4C and S6). To provide additional evidence that HK2 associates with the 5’UTR of the *SOX10* mRNA, we used an immunofluorescence-based RNA-Proximity Ligation Assay (RNA-PLA) (Zhang *et al*, 2016). The RNA-PLA assay enables the study of the association of proteins of interest with their target RNAs *in situ*. The association of HK2 with the *SOX10* mRNA was observed by a robust RNA-PLA signal (Figure 4D), which was generated by the use of an antibody that specifically recognizes HK2, in combination with an antisense probe that hybridize close to the predicted HK2-*SOX10* interaction region within the 5’UTR. To further confirm the specificity of the PLA signal, we depleted HK2 or SOX10 using target-specific siRNAs. As expected, siRNA-mediated depletion of HK2 or *SOX10* mRNA resulted in a loss of proximity signal when compared to the cells transfected with control siRNA (Figure 4D and S6C), demonstrating the specificity of the signal for HK2-*SOX10* 5’UTR interactions. Together, these results indicate that HK2 preferentially associates with *SOX10* via its 5’UTR.

### The 3’ half of the *SOX10* 5’UTR is essential for *SOX10* 5’UTR-driven translation and HK2 association

Next, we sought to define the *cis*-acting elements on the *SOX10* mRNA involved in the translational regulation by HK2. To this end, we performed a luciferase assay coupled with RT-qPCR using constructs containing the full-length (FL) 5′ and 3’ UTR of *SOX10* mRNA, which were cloned upstream and downstream of a Renilla ORF, respectively. We also generated six *SOX10* 5’UTR deletion mutants containing distinct deletions of ∼50 nucleotides each (Δ1 to Δ6, based on the region of the deletion) (Figure 5A). In this assay, we transiently transfected the Renilia reporters together with a control Firefly luciferase expressing plasmid into A375 cells and quantified the relative Renilia over Firefly luciferase activities (Rluc/Fluc ratio). We observed a higher Rluc/Fluc ratio for the construct containing the 5’UTR compared to the 3’UTR or no UTR (“empty”) showing the importance of the 5’UTR in the translation of the *SOX10* mRNA. We also observed that the constructs expressing the Δ4, Δ5 and Δ6 deletions (but not the Δ1, Δ2 and Δ3 deletions) presented a reduced Rluc/Fluc ratio when compared to the 5’UTR-containing plasmid (Figure 5B). Of note, the different *SOX10* vectors expressed a similar level of RNA (Figure 5C), suggesting that the 3’ half of the *SOX10* 5’UTR (152-301 nts) is essential for efficient translation. We next explored the possibility that the observed translation effects were mediated by HK2 association to the *SOX10* 5’UTR. Using RIP experiments, we found a reduced association between HK2 and the Δ4, Δ5 and Δ6 deletion mutants of the 5’UTR RNA (with a significant effect for the Δ4 deletion) (Figure 5D). Together, these results indicate that the 3’ half of the *SOX10* 5’UTR is crucial for HK2 association and for driving *SOX10* 5’UTR-mediated translation.

**Figure 5.**
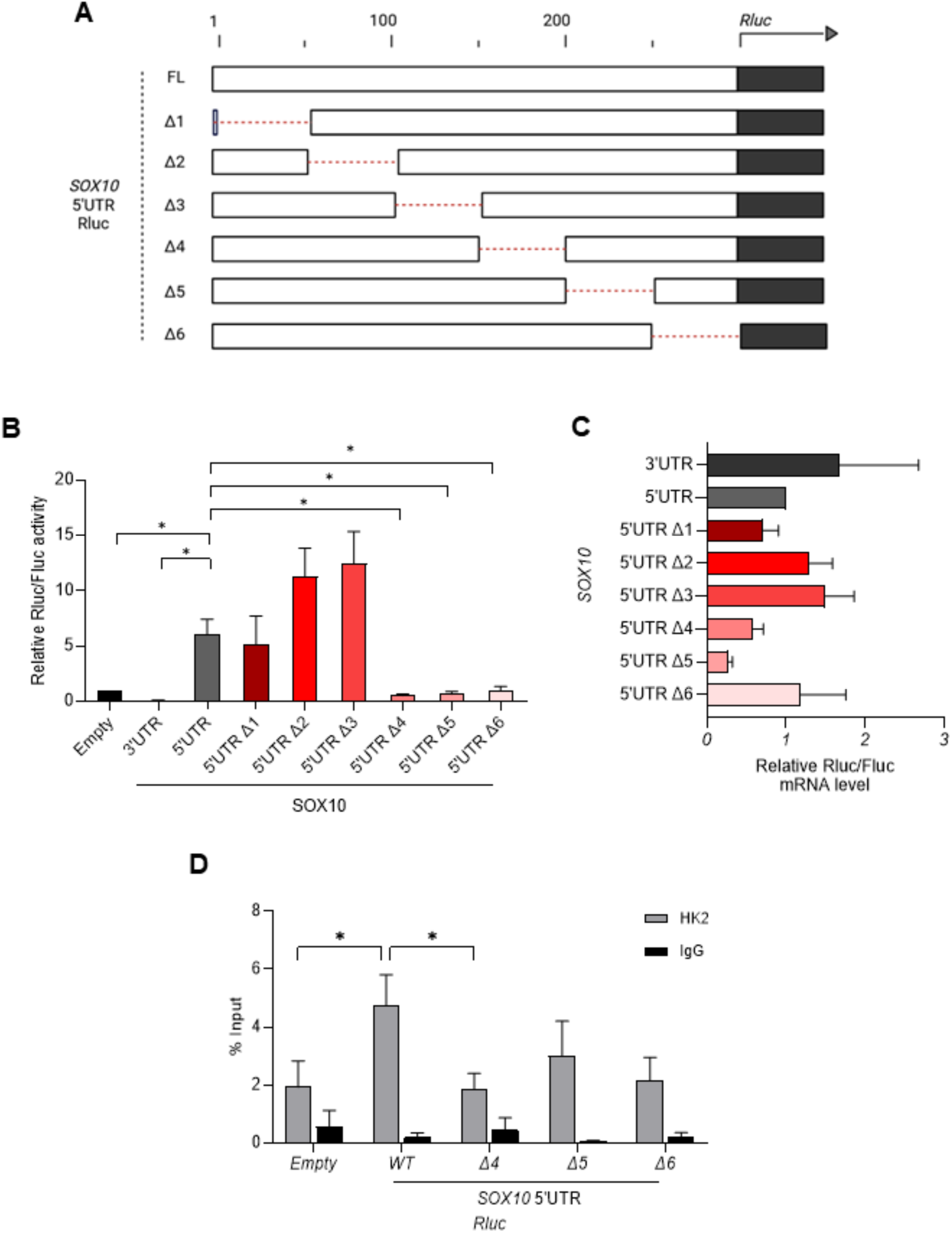
The 3’ half of the *SOX10* 5’UTR is essential for HK2 association and *SOX10* 5’UTR-driven translation. **(A)** Schematic of full-length (FL) and mutated *SOX10* 5’UTR (Δ1-6) reporters cloned upstream of the Renilla reporter. The deleted regions of ∼50nts are represented as red dashed lines. **(B)** Luciferase assay performed in A375 cells co-transfected with a Firefly reporter and the Renilla reporters in (A). An empty Renilla vector and the *SOX10* 3’UTR Renilla reporter were used as controls. The activity of the Firefly and Renilla reporters was measured 48h after transfection. The Renilla activity was normalized by the Firefly activity, and the data shown is relative to the Empty vector. *p*-values were calculated by Mann-Whitney statistical test (SEM, n = 4 biological replicates) (∗ p < 0.05). **(C)** RT-qPCR quantification of the *Renilla* mRNA from the same transfected cells in (B). The *Renilla* expression was normalized by the *Firefly* expression, and the data shown is relative to the expression of the Empty vector. *p*-values were calculated by Mann-Whitney statistical test (SEM, n = 4 biological replicates). **(D)** RIP experiment performed on A375 cells transfected with Renilla luciferase reporters containing the *SOX10* 5’UTR FL and Δ4, Δ5 and Δ6. *Renilla* mRNA was analyzed in IgG and HK2 immunoprecipitated samples and expressed as percentage of the mRNA present in the input. *p*-values were calculated by ordinary two-way ANOVA with Dunnett’s multiple comparisons test (SEM, n = 3 biological replicates) (* p ≤ 0.05).

### HK2-dependent *SOX10* translational regulation is involved in colony formation and proliferation properties of melanoma cells

We next investigated the possibility that the HK2-*SOX10* 5’UTR interaction is involved in cancer-related phenotypes. Functional assays were conducted in A375 and A2058 melanoma cell lines upon HK2 knock down. To investigate whether the effects observed by depleting HK2 are mediated by the 5’UTR-dependent translation regulation of the *SOX10* mRNA, we ectopically expressed SOX10 in both cell lines through the transduction of lentivirus particles carrying the sequence of the human SOX10 open reading frame (ORF) (Figure 6A), without the 5′UTR. We found that HK2 knock down led to decreased colony formation and proliferation of melanoma cells (Figure 6B-C). As expected, SOX10 ectopic expression reduced the effect of HK2 depletion on colony formation, while rescuing the proliferation properties of the melanoma cells (Figure 6B-C). Collectively, our findings indicate that the oncogenic functions of HK2 in melanoma cells are, at least in part, dependent on the translational regulation of the *SOX10* mRNA via its 5’ UTR.

**Figure 6.**
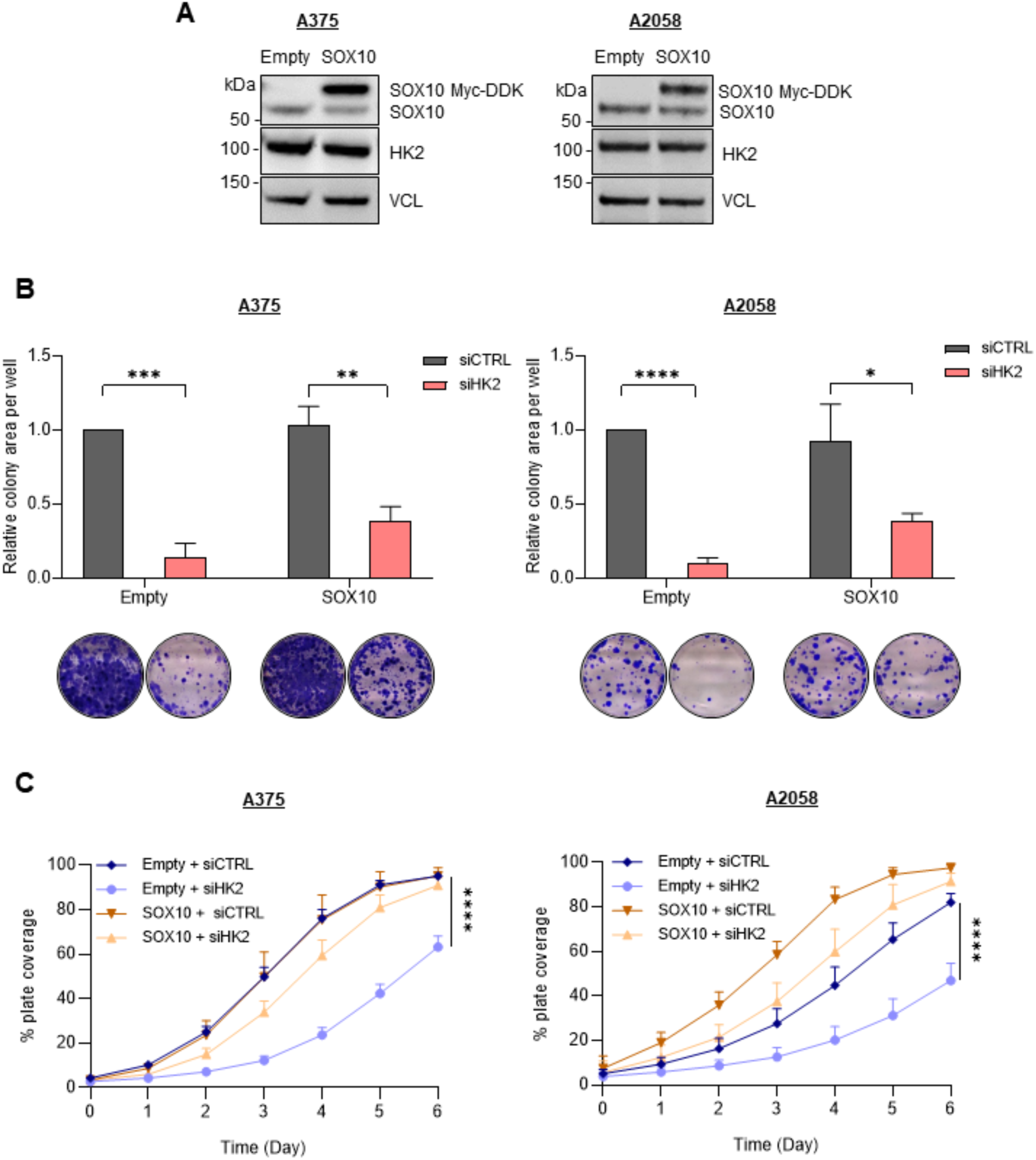
HK2-dependent SOX10 translational regulation is involved in clonogenicity and proliferation properties of melanoma cells. **(A)** Western blot analysis of the ectopic SOX10 Myc-DDK expression in A375 and A2058 melanoma cells. VCL was used as loading control (n = 3 biological replicates). **(B)** A375 and A2058 cells from (A) were plated 24h after transfection with siRNAs targeting HK2 (siHK2, light red) or control (siCTRL, grey). The colonies were stained with crystal violet after 10 days, and the clonogenic cell growth was measured. Upper panel: percentage of area covered by crystal violet stained cell colonies. Lower panel: representative images of three independent experiments. *p*-values were calculated by ordinary two-way ANOVA with Turkey’s multiple comparison test (SD, n = 3 biological replicates), and only significant differences within the same cell line are shown (* p ≤ 0.05, **** p ≤ 0.0001). **(C)** Cell proliferation assay performed in A375 and A2058 cells from (A). Cells were plated 24h after transfection with siRNAs targeting HK2 (Empty, light blue; SOX10, light orange) or control (Empty, dark blue; SOX10 dark orange). *p*-values were calculated by ordinary two-way ANOVA with Turkey’s multiple comparison test (SD, n = 3 biological replicates), and only significant differences within the same cell line are shown (**** p ≤ 0.0001).

## DISCUSSION

The glycolytic enzyme HK2 has recently been found to perform non-glycolytic activities in cancer, including regulation of transcription (Wang *et al*, 2017), anti-apoptotic (Zheng *et al*, 2023; AZOULAY-ZOHAR *et al*, 2004), scaffolding activities (Blaha *et al*, 2022), and protein kinase activities mediating tumour immune evasion (Guo *et al*, 2022). In yeast, hexokinases have been shown to directly bind to nucleic acids (Asencio *et al*, 2018), thus suggesting unexpected DNA- or RNA-binding activities. Although HK2 does not harbour any recognisable RNA-binding domain, this kinase has been proposed to bind to poly(A) and non-poly(A) RNAs in human cells in large-scale RNA interactome studies (Backlund *et al*, 2020; Trendel *et al*, 2019; Queiroz *et al*, 2019; Urdaneta *et al*, 2019). In this work, we demonstrated that HK2 is a novel *bona-fide* RNA-binding protein implicated in the control of mRNA translation in melanoma. We found that HK2 specifically binds the *SOX10* mRNA, a key transcriptional factor involved in tumour initiation, and regulates mRNA translation initiation in a 5’UTR-dependent manner. We found that the *SOX10* 5’UTR is essential for HK2 association and translation.

Some reports have demonstrated the involvement of the RNA-binding activity exerted by GAPDH in the control of mRNA stability through interaction with 3’UTRs of several mRNAs in cancer cells (Kondo *et al*, 2011; Bonafé *et al*, 2005; Zhou *et al*, 2008; Ikeda *et al*, 2012). In addition, PKM has been shown to promote ribosome stalling in the vicinity of glutamate- and lysine-encoding regions in ORFs, thereby inhibiting translation elongation (Kejiou *et al*, 2023). Our findings extend these studies, providing new evidence that the moonlighting RNA-binding activity of glycolytic enzymes (*i.e.* HK2) plays a key role in translational initiation through an interaction with a 5’UTR.

Since (i) regulation of translation initiation is a crucial mechanism for gene expression, dynamically regulating protein synthesis and thereby contributing to the determination of the cellular phenotype (Song *et al*, 2021; Fabbri *et al*, 2021), and (ii) our results demonstrate that the HK2-*SOX10* 5’UTR RNA-protein interaction is involved in cancer-relevant processes such as the ability of melanoma cells to proliferate and form colonies, further research is needed to unravel the factors involved in the regulation of *SOX10* mRNA translation by HK2. This is particularly important since HK2 is an attractive target in cancer (Shan *et al*, 2022) and since SOX10 is not only critical in melanoma proliferation but also a factor of resistance to immunologic cell death (Rosenbaum *et al*, 2024).

Our findings underscore a non-metabolic function of HK2, indicating that cancer cells may enhance glycolysis for purposes beyond simple anabolism. While aerobic glycolysis provides significant advantages for rapidly growing and proliferating cells, contributing to 60% of total cellular ATP production in cancer cells (Busk *et al*, 2008) and supplying glycolytic intermediates necessary for the biosynthesis of nucleic acids, lipids, and amino acids (Vander Heiden *et al*, 2009), its role extends further. Given that many glycolytic enzymes also serve as RNA-binding proteins engaged in post-transcriptional RNA-processing reactions and considering that aberrant mRNA translation is a hallmark of tumours, it is crucial to investigate the significance of HK2 interactions with RNA beyond the *SOX10* mRNA. HK2 might regulate the translation of several other key mRNAs involved in cancer-relevant signalling pathways. Understanding the cellular processes influenced by HK2 RNA-binding activity and uncovering their underlying molecular mechanisms represent promising directions for future research. Such studies could lead to the development of novel therapeutics that specifically block HK2-RNA interaction in cancer cells while sparing non-malignant counterparts, offering new perspectives in cancer treatment.

## METHODS

### Cell culture, transfection and transduction

M249 cells were grown in RPMI 1640 (Biowest, #L0500) supplemented with 10% FBS (Gibco™, #A5256701) and 2 mM L-glutamine (Eurobio-Scientific, #CSTGLU00-0U). MALME 3M cells were grown in high glucose (4.5 g/L) DMEM supplemented with 20% FBS. HEK293T, A375, A258 and other melanoma cells were grown in high glucose DMEM (Biowest, #L0101-500) supplemented with 10% FBS and 2 mM L-glutamine. All cells were grown in 5% CO2 humidified incubator at 37°C. When indicated, the high glucose medium of A375 cells was replaced with glucose-free DMEM (Gibco™, #11966025) containing 10% FBS and 2 mM L-glutamine and supplemented with 5 g/L galactose (Sigma-Aldrich, #G5388) or 16mM 2-DG (MedChemExpress, #HY-13966) when indicated. The cells were washed with PBS before the medium was replaced. Cells were regularly tested for mycoplasma and authenticated by short tandem repeat (STR) profiling.

Cells were plated one day before for siRNA and plasmid transfections. The siRNAs were purchased from Dharmacon (ON-TARGETplus technology), and siRNA transfections were performed using Lipofectamine RNAiMAX (Invitrogen™, #13778150) according to manufacturer’s instructions. Transfected cells were either harvested at 48h or re-plated 24 h after transfection, as indicated in the figure legends. Plasmid transfections were performed using Lipofectamine 2000 (Invitrogen™, #11668019) or JetPEI (Polyplus Transfection, #101000020), and the transfection was performed according to manufacturer’s instructions. At 48 h post-transfection, cells were harvested for further analysis. Lipofectamine 2000 was used for the transfection of Renilla and Firefly luciferase reporters in A375 cells. The cDNA for human *SOX10* 5’UTR and 3’UTR were subcloned into the pRP(Exp)-EGFP-CMV vector upstream and downstream a Renilla sequence, respectively, and they were kindly provided by Prof Caroline Robert (Gustave Roussy, France). Site-directed mutagenesis of *SOX10* 5’UTR was performed using Q5® Site-Directed Mutagenesis Kit (New England Biolabs, #E0554) according to the manufacturer’s instructions. JetPEI was used for transient transfection of HK2 plasmids in HEK293T cells. HK2 (NM_000189.5) plasmids were purchased from VectorBuilder. HK2-WT (wild-type; #VB220927-1092xyz), DA (D209A/D657A, a non-glucose-binding mutant; #VB221214-1257apk), SA (S340A/S788A, a catalytically inactive mutant; #VB221214-1258ubs), and MBD (deletion of 1–15aa, mitochondrial-binding deficient; #VB221214-1255ezg) were cloned into a pLV[Exp]-Puro-CMV EGFP expressing vector.

To establish the stable overexpression of SOX10 (Myc-DDK tagged; OriGene, #RC203545L3V) in A375 and A2058 cell lines, cells were transduced with pLenti-C-Myc-DDK-P2A-Puro vector expressing SOX10 (NM_006941) ORF nucleotide sequence (OriGene, #RC203545L3V) according to manufacturer’s instructions. Lentivirus infection was performed during 48h, and cells were selected with puromycin (2 μg/ml). Western blot and RT-qPCR analyses were performed to confirm the ectopic expression of SOX10 in the stable cells. The stablished cell lines were cultured in high glucose DMEM medium with 10% FBS and 2 mM L-glutamine supplemented with puromycin (2 μg/ml).

### Polysome profiling

Polysome profiling was performed as previously described (Shen *et al*, 2019; Boussemart *et al*, 2014). Briefly, A375 cells were incubated with 100 μg/mL cycloheximide in fresh medium at 37 °C for 5 min. Cells were then washed, scraped into ice-cold PBS supplemented with 100 μg/mL cycloheximide (Sigma-Aldrich, #C4859), and centrifuged for 10 min at 3000 rpm. The cell pellets were re-suspended into 400 μL of LSB buffer (20 mM Tris, pH 7.4, 100 mM NaCl, 3 mM MgCl2, 0.5 M sucrose, 1 mM DTT, 100 U/mL RNasOUT and 100 μg/mL cylcoheximide). After homogenization, 400 μL LSB buffer supplemented with 0.2 % Triton X-100 and 0.25 M sucrose was added. Samples were incubated on ice for 30 min, and then centrifuged at 12,000 × g for 15 min at 4 °C. The supernatant was adjusted to 5 M NaCl (Sigma-Aldrich, #S6546) and 1 M MgCl2 (Invitrogen™, AM9530G). The lysates were then loaded onto a 5–50% sucrose density gradient and centrifuged in an SW41 Ti rotor (Beckman) at 36,000 rpm for 2 h at 4 °C. Polysome fractions were monitored and collected using a gradient fractionation system (Isco). The sucrose gradient fractions were stored at −80°C or directly processed for western blot analysis or RNA extraction. For RT-qPCR analysis, RNAs from each fraction were extracted using TRIzol-LS (Invitrogen™, #10296028) according to manufacturer’s procedure and were quantified using RNA 2100 Bioanalyzer (Agilent Genomics). The expression of a panel of 84 EMT-associates genes was measured using RT^2^ Profiler PCR Array (qPCR Qiagen EMT array, #330231) in both total RNA and polysomal RNA levels of A375 cells transfected with siRNAs targeting HK2 or control.

### RNA sequencing and bioinformatic analysis

RNA sequencing libraries were prepared from 500ng to 1µg of total RNA or mRNA-enriched from heavy polysome fractions using the Illumina TruSeq Stranded mRNA Library preparation kit which allows to perform a strand specific sequencing. Nanodrop spectrophotometer was used to assess purity of RNA based on absorbance ratios (260/280 and 260/230) and BioAnalyzer for RNA integrity (RIN>9). A first step of polyA+ selection using magnetic beads is done to focus sequencing on polyadenylated transcripts. After fragmentation, cDNA synthesis was performed and resulting fragments were used for dA-tailing followed by ligation of TruSeq indexed adapters. PCR amplification was finally achieved to generate the final barcoded cDNA libraries. The libraries were equimolarly pooled and subjected to qPCR quantification using the KAPA library quantification kit (Roche). Sequencing was carried out on the NovaSeq 6000 instrument from Illumina based on a 2*100 cycle mode (paired-end reads, 100 bases) to obtain around 30 million clusters (60 million raw paired-end reads) per sample. Finally, Fastq files were generated from raw sequencing data using bcl2fastq pipeline performing data demultiplexing based on index sequences.

RNA-seq data were analyzed with the Institut Curie RNA-seq Nextflow pipeline (Servant *et al*, 2023) (v3.1.4). Briefly, raw reads were first trimmed with Trimgalor. Reads were aligned on the human reference genome (hg19) using STAR (STAR_2.6.1a_08-27). Genes abundances were then estimated using STAR, and the Gencode v34 annotation. Differential analysis between conditions were done using the R package Xtail (Xiao *et al*, 2016) only on protein coding genes.

We used the variance stabilizing transformation (VST) offered by *DESeq2* (Love *et al*, 2014) on the raw count data to stabilize the variance across the mean and then performed a principal components analysis (PCA). The PCA plots (Figure S7) were built using the ggplot2 package (Wickham, 2009).

### Complex Capture (2C)

A375 cells were irradiated with UV-C light at 254 nm and lysed in HMGN150 buffer (20 mM HEPES pH 7,5; 150 mM NaCl; 2 mM MgCl2; 0.5% Nonidet P−40; 10% Glycerol). Lysates were cleared via centrifugation at 10,000 × g for 10 minutes at 4°C, and then treated or not with RNAse A (15μL of RNAse A at 10 mg/mL for 1 mg of proteins; Thermo Scientific™, #EN0531). A fraction of the input (2%) was kept as control for the protein expression on SDS-PAGE. The Quick-RNA Midiprep kit was to purify crosslinked nucleic acids–RBP complexes. RNA concentration of the eluate was measured using NanoDrop (Thermo Scientific™), and 50 μg RNA was treated with RNAse I (500 U; Invitrogen™, #AM2295) for 40 min at 30°C. The samples were then mixed with 1X loading buffer for subsequent capillary western blot analysis.

### CrossLinking and immunoprecipitation (CLIP)

Cells were cultured on a 15 cm dish until 70-80 % of confluence. The medium was removed, and the cells were washed with ice-cold PBS and irradiated with UV-C light at 254 nm. The cells were then lysed in 1 mL RIPA lysis buffer (50 mM Tris-HCl pH 7.4, 100 mM NaCl, 1% Nonidet P-40, 0.1% SDS, 0.5% Sodium deoxycholate), supplemented with RNAseOUT (40 U/mL; Invitrogen™, #10777019) and protease inhibitor cocktail (complete EDTA free; Roche, #11873580001). Lysates were cleared via centrifugation at 18,000 x g for 10 minutes at 4°C, and then treated or not with RNAse I (0.3 – 1 U) in combination with TURBO™ DNase I (4 U; Invitrogen, #AM2238) and incubated at 37°C for 3 minutes at 800 rpm. 1% of the lysate was used as input material to quantify total protein concentration in western blot analysis. The rest of the lysate was incubated with the HK2 antibody-conjugated beads or GFP-Trap® Magnetic Agarose (ChromoTek, #gtma) overnight at 4°C while constantly rotating. For endogenous HK2 immunoprecipitation, 9 µg of HK2 antibody (Abcam, ab209847) or appropriate control IgG (rabbit; Cell Signaling, #2729S) were coupled overnight at 4°C to 90µl Dynabeads Protein G (Invitrogen™) while constantly rotating, and prior to cell harvesting. Conversely, GFP-Trap® Magnetic Agarose was used for the immunoprecipitation of HK2 mutants conjugated with GFP. The immunocomplexes were washed twice with RIPA-S buffer (50 mM Tris- HCl pH 7.4, 1 M NaCl, 1 mM EDTA, 1% Nonidet P-40, 0.1% SDS, 0.5% Sodium deoxycholate) and twice with PNK buffer (20 mM Tris-HCl pH 7.4, 10 mM MgCl_2_, 0.2% Tween 20). After washes, 10% of the immunocomplexes was used to validate the correct immunoprecipitation by western blot analysis. Then, crosslinked nucleic acids were radiolabeled with ATP, [γ- 32P] (PerkinElmer) by T4 polynucleotide kinase (PNK; New England Biolabs, #M0201S) for 30 minutes at 37°C. After two washes with RIPA-S buffer and one wash with PNK buffer, protein-RNA complexes were eluted in 1X loading buffer at 75°C for 10 minutes and separated by SDS-PAGE. Then the gel was dried, and complexes were exposed to a phosphorimaging screen and scanned with Amersham™ Typhoon™ Biomolecular Imager (CYTIVA). The files were processed with ImageJ software.

### Sucrose density gradient ultracentrifugation and fractionation

For the sucrose density gradient ultracentrifugation and fractionation assay, previously published protocols were used as a basis (Huppertz *et al*, 2022; Caudron-Herger *et al*, 2019). A375 cells were cultured on a 15 cm dish and lysed in 300 μl lysis buffer (25 mM Tris-HCl pH 7.4, 2 mM EDTA, 150 mM KCl, 1 mM NaF, 0.5 mM DTT, 0.5% Nonidet P-40), supplemented with protease inhibitor cocktail (complete EDTA free; Roche, #11873580001). As control, lysis buffer was also supplemented with RNAseOUT (40 U/mL; Invitrogen™, #10777019). Lysates were pre-cleared via centrifugation at 10,000 × g for 10 minutes at 4°C. The lysates were then treated with a combination of RNase I (150 U; Invitrogen™, #AM2295) and RNase A/T1 (50 U / 125 U; Thermo Scientific™, #EN0551) and incubated at 37°C for 15 minutes. For the control sample, RNaseOUT was added to the lysate and the sample was incubated for 15 min at 4°C. For the fractionation, sucrose gradients were prepared from 5% (w/v) to 25% (w/v) sucrose (in 150 mM KCl, 25 mM Tris-HCl pH 7.4 and 2 mM EDTA) using Isco Model 160 Gradient Former. Lysates were then separated by centrifugation at 30,000 rpm and 4°C in an SW41Ti rotor (Beckman) for 18 hours. After ultracentrifugation, the lysate fractions were monitored and carefully collected using a gradient fractionation system (Isco) (20 fractions were collected of approximately 1000 μl) and transferred into fresh 1.5 mL tubes. For the protein precipitation, 150 μl of cold 100% Trichloroacetic acid (TCA; Sigma-Aldrich, #T6399) was added, and the samples were incubated on ice for 30 minutes. The fractions were centrifuged at 18,000 x g for 20 minutes at 4°C. The TCA supernatant was carefully removed, and the pellets were washed once with 1 ml cold acetone (stored at −20°C). The fractions were vortexed, and an additional centrifugation step was performed at 18,000 x g for 30 minutes at 4°C. The supernatant was carefully removed, and the pellet air-dried. Finally, the pellets were resuspended in 20 μl 1X loading buffer, incubated at 95°C for 5 minutes, and used for SDS-PAGE and immunoblotting.

### RNA immunoprecipitation (RIP) assay

Prior to harvesting cells, 9 µg of antibody anti-HK2 (Abcam, ab209847) were coupled overnight at 4°C to 90µl Dynabeads Protein G (Invitrogen™) while constantly rotating. A375 and A2058 cells were cultured on a 15 cm dish and lysed in 500 μl Co-IP buffer (50 mM Tris-HCl pH 7.4, 1 mM EDTA, 150 mM NaCl, 0.1% Nonidet P-40), supplemented with protease inhibitor cocktail (complete EDTA free; Roche, #11873580001) and RNAseOUT (40 U/mL; Invitrogen™, #10777019) for 30 minutes at 4°C. Lysates were cleared via centrifugation at 18,000 x g for 10 minutes at 4°C. After lysis, 1% and 10% of the lysate was used as input material to confirm total protein and RNA levels, respectively. The rest of the lysate was incubated with the HK2 antibody-conjugated beads overnight at 4°C while constantly rotating. The immunocomplexes were washed five times with ice-cold CO-IP buffer supplemented with protease inhibitor cocktail and RNAseOUT. After washes, 10% of the immunocomplexes was used to validate the correct immunoprecipitation by western blot analysis. The rest of the beads were eluted with 100 μl Protein-RNA elution buffer (100 mM Tris-HCl pH 8, 10 mM EDTA, 1% SDS) at 80°C for 5 minutes, followed by another 5 minutes incubation at room temperature. To recover the RNA bound by HK2, the beads were resuspended in TRIzol (Invitrogen™, #15596018), as well as the input RNA, and RNA extraction was followed according to manufacturer’s procedure. The extracted RNA was then used for subsequent RT-qPCR experiments.

### *In silico* binding prediction

The catRAPID algorithm (Armaos *et al*, 2021) was used to estimate the potential binding sites of HK2 on *SOX10* mRNA. The highest-raking site at RNA position 206-321 bp was used for subsequent analysis.

### Seahorse metabolic flux assay

Oxygen consumption rates (OCR) and extracellular acidification rates (ECAR) were analysed on a XF96 Extracellular Flux Analyzer (Seahorse Bioscience). A375 cells were plated in non-buffered DMEM medium supplemented with 25 mM glucose (Sigma-Aldrich, #D9434) or galactose (Sigma-Aldrich, #G5388). Measurements were obtained under basal conditions (no treatment) and after the addition of mitochondrial inhibitors (oligomycin, 1μM; FCCP, 0.5μM; rotenone/antimycin A, 0.5μM), or glycolysis inhibitor (2-DG, 16mM) (MedChemExpress, #HY-13966).

### Colony formation assay

Cells transfected with 25nM siRNA (Dharmacon) were seeded in triplicates in six-well plates. After 10 days of incubation, cells were washed with PBS 1X (Gibco™, #70011044) and stained with 0.5% of crystal violet (Sigma Aldrich, #C0775) for 15 min. Plates were washed with water and dried before imaging. Colony area was quantified using ImageJ-plugin “ColonyArea”(Guzmán *et al*, 2014).

### Luciferase assay

A375 cells were transiently transfected with Firefly and Renilla luciferase reporters using Lipofectamine 2000 Reagent (Invitrogen™, #11668019). Renilla vectors containing the full-length (FL) *SOX10* 3’UTR or 5′UTR, as well as six *SOX10* 5’UTR mutants (Δ1-Δ6) harboring deletions of ∼50 nucleotides, were transfected in the cells. An empty Renilla vector (without *SOX10* 5’ or 3’UTR) was used as control for the experiment. After 48h, part of the transfected cells was harvested for RNA extraction to measure the vectors’ RNA expression by qPCR, while the rest of the cells were used to measure the luciferase activity using the Dual-Luciferase Reporter Assay System (Promega). Data was presented as the ratio between Renilla and Firefly luciferase RNA expression or luminescence activities, respectively.

### Proliferation assay

Cells transfected with 25nM siRNA (Dharmacon) were seeded in triplicates in six-well plates. The cell growth was analyzed for percent plate coverage with IncuCyte S3 Live Cell Analysis System (Sartorius). Pictures of each well were acquired every 3 hours for 7 days.

### Protein extracts, SDS-PAGE and Western blot

Whole cell lysates were prepared using RIPA buffer supplemented with RNaseOUT (40 U/mL; Invitrogen™, #10777019) and protease inhibitor cocktail (complete EDTA free; Roche, #11873580001). Lysates were cleared via centrifugation at 18,000 x g for 10 minutes at 4°C. Cell lysates were quantified for protein concentration using a bicinchoninic acid (BCA) protein assay kit (Thermo Scientific™, #23225). Protein samples were resolved on NuPAGE 4–12% Bis-Tris gels with MOPS buffer or 3–8% Tris-acetate gels with Tris-acetate buffer (Life Technologies, #LA0041) and then transferred to 0.45 μm nitrocellulose membrane (Amersham). After saturation in Tris-buffered saline buffer containing 0.01% Tween-20 (TBST) and supplemented with 5% powdered milk, the membranes were incubated with primary antibodies either for 1 hour at room temperature or overnight at 4 °C with agitations, followed by 3 X TBST washes of 15 minutes each, and incubation of the membranes with peroxidase (HRP)- conjugated secondary antibodies diluted in 5% milk TBST for one hour at room temperature. After 3 X TBST washes of 5 minutes each, the membranes were incubated in ECL solution for development.

The following antibodies were used for assays: anti-HK2 (Abcam, #ab209847; 1:1,000 dilution), anti-SOX10 (Santa Cruz Biotechnology, #sc365692; 1:1,000 dilution), anti-GFP (Roche, #11814460001; 1:1,000 dilution), anti-HUR (Santa Cruz Biotechnology, #sc-5261; 1:1,000 dilution), anti-H3 (Abcam, #ab1791; 1:500 dilution), anti-GAPDH (Sigma-Aldrich, #G9545; 1:5,000), anti-PKM2 (Cell Signaling, #4053S; 1:1,000 dilution), anti-LDHB (Santa Cruz Biotechnology, #sc100775; 1:1,000 dilution), anti-S6 ribosomal protein (Cell Signaling, #2317; 1:1,000 dilution), anti-VCL (Abcam, #ab129002; 1:3,000 dilution), anti-alpha Tubulin antibody (GeneTex, # GTX628802; 1:1000 dilution), anti-HSC70 antibody (Santa Cruz Biotechnology, #sc-7298; 1:1,000 dilution), anti-ACOT1/2 (Santa Cruz Biotechnology, #sc-373917; 1:1000 dilution), goat anti-rabbit (Sigma-Aldrich, Thermo Fisher Scientific, 1:2,000 dilution) and goat anti-mouse (Sigma-Aldrich, Thermo Fisher Scientific, 1:5,000 dilution) antibodies.

### RNA extraction and RT-qPCR

RNA was isolated using the TRIzol-chloroform method and treated with DNase I (TURBO DNA-free; Invitrogen™, #AM1907) according to the manufacturer’s instructions. After DNase treatment, RNA was quantified using a Nanodrop 2000 spectrophotometer (Thermo Scientific™), and 1 μg total RNA was used to synthesize cDNA using SuperScript III Reverse Transcriptase (200 U/μL; Invitrogen™, #18080044) and random hexamer primers (Invitrogen™, #SO142), according to the manufacturer’s instructions. Quantitative PCR (qPCR) was performed using the Power SYBR Green PCR Master Mix (Thermo Scientific™, #4367559) on a CFX96 Real-Time PCR Detection System (Bio-Rad Laboratories Inc). Primer sequences are given in Table S3. Gene expression values were determined relative to VCL and are shown as a relative fold change to the value of control samples. All experiments were performed in biological triplicates and error bars indicate ± standard deviation as assayed by the ΔΔCt method.

For RT-qPCR analysis of polysome fractions, RNA of each fraction (250 μL), comprising both monosome fractions and polysome fractions, was extracted using TRIzol LS (Invitrogen™, #10296028). The extracted RNA was diluted in 30 μL RNase-free water (Invitrogen™, #10977035). For each fraction, the same volume of RNA was retrotranscribed into cDNA using SuperScript IV Reverse Transcriptase (Invitrogen™, #18090010) and random hexamer primers (Invitrogen™, #SO142), according to the manufacturer’s instructions. Following the reverse transcription, mRNA abundance was determined by qPCR using Power SYBR Green PCR Master Mix (Thermo Scientific™, #4367559). Data were analyzed by the threshold cycle (Ct) comparative method and quantified as percentage of the total RNA considering the whole fractions stand for 100%. HPRT gene was used as a control.

### RNA Proximity Ligation Assay (RNA-PLA)

Interactions between HK2 and SOX10 mRNA were monitored *in cellulo* by RNA-PLA (Zhang *et al*, 2016). The experiment was performed using Duolink In situ Detection Reagents FarRed according to the manufacturer’s instructions (Sigma-Aldrich, #DUO92013). A375 cells were seeded on coverslips 45mn 24 h before transfection with siRNAs targeting HK2 (siHK2), SOX10 (siSOX10) or control (siCTRL). After 48 h, cells were fixed with 4% PFA in PBS for 30 minutes at room temperature. Next, coverslips were washed with PBS and permeabilized with Tween 0.2% in PBS for 10 minutes at room temperature. Cells were then washed with PBS and blocked using a blocking buffer (10% goat serum and 0.1% Triton X-100 in PBS supplemented with 10 ug of salmon sperm) for 1 h at room temperature. Cells were once again washed with PBS and incubated in blocking buffer containing 100 nM of an anti-sense PLUS probe in a wet chamber at 4°C overnight. Salmon sperm was boiled for 3 minutes at 70°C prior to addition. Of note, this probe was composed of a 20mer of DNA complementary to the SOX10 5’UTR, a polyA linker and a sequence complementary to the PLA MINUS probe: GCGGTCCAGCTCGGGGCTGGGAGGTGACGCTGGTGGGCTGGGAGGGAAAAAAAAAAAAAAAAAAAAATATGACAGAACTAGACACTCTT.

Subsequently, cells were washed with PBS and incubated in blocking buffer for 1 h at room temperature. Then, HK2 primary antibody (Abcam, # ab209847) was diluted in blocking buffer (1.1000) and incubated for 1 h at 37°C. Next, the Duolink® PLA® probes anti-Rabbit MINUS (5X) containing secondary antibodies conjugated to oligonucleotides were added, and the mixture was incubated for 1 hour at 37°C. A ligation solution is then added to ligate the complementary oligonucleotides of the two probes (PLUS and MINUS) if in proximity (< 40 nm) to form a closed circle by incubating the coverslips 45mn at 37°C. This DNA was subsequently amplified by rolling circle replication: Duolink polymerase and its buffer are spread on the coverslips, then incubated for 2 hours at 37°C. The coverslips were finally mounted in ProLong™ Diamond Antifade Mountant with DAPI (Invitrogen™, #P36935) to visualize the nuclei and the amplicons were detected by far-red fluorescent probes. Images of each coverslip were acquired using Leica 3D SIM (Structured Illumination Microscopy) microscope at the PICT-IBiSA Imaging Facility in Orsay, and PLA spots were analyzed using the semi-automatic macro Mic-Maq on Fiji software.

### Quantification and statistical analysis

Statistical analyses were performed on GraphPad Prism (Version 10.3.0) and statistical significance defined as a P value < 0.05 was determined by two-tailed unpaired Student’s t test. Comparisons in multiple groups were analyzed with one-way or two-way ANOVA, as indicated in the figure legends. All data in this study are presented as the mean standard deviation or mean standard error.

## ACKNOWLEDGEMENTS

This work was supported by grants from Institut Curie, Gustave Roussy, INSERM, CNRS, Equipe labellisée Ligue Nationale Contre le Cancer (LNCC) (to SV), La fondation Carrefour, la Fondation du Crédit Mutuel, et Sébastien Bazin (to CR). This project has received funding from the European Union’s Horizon 2020 research and innovation program under the Marie Skłodowska-Curie grant agreement No 847718. High-throughput sequencing was performed by the ICGex NGS platform of the Institut Curie supported by the grants ANR-10-EQPX-03 (Equipex) and ANR-10-INBS-09-08 (France Génomique Consortium) from the Agence Nationale de la Recherche (“Investissements d’Avenir” program), by the ITMO-Cancer Aviesan (Plan Cancer III) and by the SiRIC-Curie program (SiRIC Grant INCa-DGOS-465 and INCa-DGOS-Inserm_12554). Data management, quality control and primary analysis were performed by the Bioinformatics platform of the Institut Curie. ALD was successively supported by pre-doctoral fellowships from the Marie Skłodowska-Curie grant agreement No 847718 and FRM (Fondation pour la Recherche Medicale). GC was supported by the Marie Sklodowska-Curie Campus France Fellowship PRESTIGE-2017-3-0017.

## AUTHOR INFORMATION

ALD performed CLIP, RIP, WB, RT-qPCR, luciferase, proliferation and clonogenic assays. AMP and GC performed polysome profiling experiments. GC and VQ performed the 2C experiment. CML performed bioinformatic analysis. DB performed the RNA-PLA experiment. RNA sequencing was performed by VR and SB. ALD, AMP, GC, CR and SV conceived the experiments and analyzed data. SV and CR coordinated the whole study. ALD and SV wrote the manuscript with the help of CR and AMP. Funding acquisition: SV, CR

## DATA AVAILABILITY

All data are available in the main text or the supplementary materials.

The datasets generated in this study have been deposited in the Gene Expression Omnibus repository (GEO) under the accession number GSE274146 (token: **qpkhimgaffylzsd**).

## SUPPLEMENTARY INFORMATION

**Suppementary Figures S1-S7**

**Suppementary Table S1**

**Suppementary Table S2 and S3**

## REFERENCES

Armaos A, Colantoni A, Proietti G, Rupert J & Tartaglia GG (2021) catRAPID omics v2.0: going deeper and wider in the prediction of protein–RNA interactions. Nucleic Acids Research 49: W72–W79

Asencio C, Chatterjee A & Hentze MW (2018) Silica-based solid-phase extraction of cross-linked nucleic acid–bound proteins. Life Science Alliance 1

Azoulay-Zohar H, Israelson A, Abu-Hamad S & Shoshan-Barmatz V (2004) In self-defence: hexokinase promotes voltage-dependent anion channel closure and prevents mitochondria-mediated apoptotic cell death. Biochemical Journal 377: 347–355

Backlund M, Stein F, Rettel M, Schwarzl T, Perez-Perri JI, Brosig A, Zhou Y, Neu-Yilik G, Hentze MW & Kulozik AE (2020) Plasticity of nuclear and cytoplasmic stress responses of RNA-binding proteins. Nucleic Acids Research 48: 4725–4740

Blaha CS, Ramakrishnan G, Jeon S-M, Nogueira V, Rho H, Kang S, Bhaskar P, Terry AR, Aissa AF, Frolov MV, et al (2022) A non-catalytic scaffolding activity of hexokinase 2 contributes to EMT and metastasis. Nat Commun 13: 899

Bonafé N, Gilmore-Hebert M, Folk NL, Azodi M, Zhou Y & Chambers SK (2005) Glyceraldehyde-3-Phosphate Dehydrogenase Binds to the AU-Rich 3′ Untranslated Region of Colony-Stimulating Factor–1 (CSF-1) Messenger RNA in Human Ovarian Cancer Cells: Possible Role in CSF-1 Posttranscriptional Regulation and Tumor Phenotype. Cancer Research 65: 3762–3771

Boussemart L, Malka-Mahieu H, Girault I, Allard D, Hemmingsson O, Tomasic G, Thomas M, Basmadjian C, Ribeiro N, Thuaud F, et al (2014) eIF4F is a nexus of resistance to anti-BRAF and anti-MEK cancer therapies. Nature 513: 105–109

Busk M, Horsman MR, Kristjansen PEG, van der Kogel AJ, Bussink J & Overgaard J (2008) Aerobic glycolysis in cancers: Implications for the usability of oxygen-responsive genes and fluorodeoxyglucose-PET as markers of tissue hypoxia. International Journal of Cancer 122: 2726–2734

Castello A, Fischer B, Frese CK, Horos R, Alleaume A-M, Foehr S, Curk T, Krijgsveld J & Hentze MW (2016) Comprehensive Identification of RNA-Binding Domains in Human Cells. Molecular Cell 63: 696–710

Castello A, Hentze MW & Preiss T (2015) Metabolic Enzymes Enjoying New Partnerships as RNA-Binding Proteins. Trends Endocrinol Metab 26: 746–757

Caudron-Herger M, Rusin SF, Adamo ME, Seiler J, Schmid VK, Barreau E, Kettenbach AN & Diederichs S (2019) R-DeeP: Proteome-wide and Quantitative Identification of RNA-Dependent Proteins by Density Gradient Ultracentrifugation. Molecular Cell 75: 184–199.e10

Chang C-H, Curtis JD, Maggi LB, Faubert B, Villarino AV, O’Sullivan D, Huang SC-C, van der Windt GJW, Blagih J, Qiu J, et al (2013) Posttranscriptional control of T cell effector function by aerobic glycolysis. Cell 153: 1239–1251

Cronin JC, Loftus SK, Baxter LL, Swatkoski S, Gucek M & Pavan WJ (2018) Identification and functional analysis of SOX10 phosphorylation sites in melanoma. PLoS One 13: e0190834

DeBerardinis RJ & Chandel NS (2020) We need to talk about the Warburg effect. Nat Metab 2: 127–129

Fabbri L, Chakraborty A, Robert C & Vagner S (2021) The plasticity of mRNA translation during cancer progression and therapy resistance. Nat Rev Cancer 21: 558–577

Graf SA, Busch C, Bosserhoff A-K, Besch R & Berking C (2014) SOX10 promotes melanoma cell invasion by regulating melanoma inhibitory activity. J Invest Dermatol 134: 2212–2220

Guo D, Tong Y, Jiang X, Meng Y, Jiang H, Du L, Wu Q, Li S, Luo S, Li M, et al (2022) Aerobic glycolysis promotes tumor immune evasion by hexokinase2-mediated phosphorylation of IκBα. Cell Metabolism 34: 1312–1324.e6

Guzmán C, Bagga M, Kaur A, Westermarck J & Abankwa D (2014) ColonyArea: An ImageJ Plugin to Automatically Quantify Colony Formation in Clonogenic Assays. PLOS ONE 9: e92444

Hafner M, Katsantoni M, Köster T, Marks J, Mukherjee J, Staiger D, Ule J & Zavolan M (2021) CLIP and complementary methods. Nat Rev Methods Primers 1: 1–23

Han S, Ren Y, He W, Liu H, Zhi Z, Zhu X, Yang T, Rong Y, Ma B, Purwin TJ, et al (2018) ERK-mediated phosphorylation regulates SOX10 sumoylation and targets expression in mutant BRAF melanoma. Nat Commun 9: 28

Huppertz I, Perez-Perri JI, Mantas P, Sekaran T, Schwarzl T, Russo F, Ferring-Appel D, Koskova Z, Dimitrova-Paternoga L, Kafkia E, et al (2022) Riboregulation of Enolase 1 activity controls glycolysis and embryonic stem cell differentiation. Molecular Cell 82: 2666–2680.e11

Ikeda Y, Yamaji R, Irie K, Kioka N & Murakami A (2012) Glyceraldehyde-3-phosphate dehydrogenase regulates cyclooxygenase-2 expression by targeting mRNA stability. Archives of Biochemistry and Biophysics 528: 141–147

Kejiou NS, Ilan L, Aigner S, Luo E, Tonn T, Ozadam H, Lee M, Cole GB, Rabano I, Rajakulendran N, et al (2023) Pyruvate Kinase M (PKM) binds ribosomes in a poly-ADP ribosylation dependent manner to induce translational stalling. Nucleic Acids Research 51: 6461–6478

Kondo S, Kubota S, Mukudai Y, Nishida T, Yoshihama Y, Shirota T, Shintani S & Takigawa M (2011) Binding of glyceraldehyde-3-phosphate dehydrogenase to the *cis*-acting element of structure-anchored repression in *ccn2* mRNA. Biochemical and Biophysical Research Communications 405: 382–387

Kudryavtseva AV, Fedorova MS, Zhavoronkov A, Moskalev AA, Zasedatelev AS, Dmitriev AA, Sadritdinova AF, Karpova IY, Nyushko KM, Kalinin DV, et al (2016) Effect of lentivirus-mediated shRNA inactivation of HK1, HK2, and HK3 genes in colorectal cancer and melanoma cells. BMC Genet 17: 156

Lee FCY & Ule J (2018) Advances in CLIP Technologies for Studies of Protein-RNA Interactions. Mol Cell 69: 354–369

Liu Z, Jia X, Duan Y, Xiao H, Sundqvist K-G, Permert J & Wang F (2013) Excess glucose induces hypoxia-inducible factor-1α in pancreatic cancer cells and stimulates glucose metabolism and cell migration. Cancer Biol Ther 14: 428–435

Love MI, Huber W & Anders S (2014) Moderated estimation of fold change and dispersion for RNA-seq data with DESeq2. Genome Biology 15: 550

Lunt SY & Vander Heiden MG (2011) Aerobic glycolysis: meeting the metabolic requirements of cell proliferation. Annu Rev Cell Dev Biol 27: 441–464

Miao W, Porter DF, Lopez-Pajares V, Siprashvili Z, Meyers RM, Bai Y, Nguyen DT, Ko LA, Zarnegar BJ, Ferguson ID, et al (2023) Glucose dissociates DDX21 dimers to regulate mRNA splicing and tissue differentiation. Cell 186: 80–97.e26

Nagy E & Rigby WFC (1995) Glyceraldehyde-3-phosphate Dehydrogenase Selectively Binds AU-rich RNA in the NAD+-binding Region (Rossmann Fold) (∗). Journal of Biological Chemistry 270: 2755–2763

Nawaz MH, Ferreira JC, Nedyalkova L, Zhu H, Carrasco-López C, Kirmizialtin S & Rabeh WM (2018) The catalytic inactivation of the N-half of human hexokinase 2 and structural and biochemical characterization of its mitochondrial conformation. Biosci Rep 38: BSR20171666

Parmenter TJ, Kleinschmidt M, Kinross KM, Bond ST, Li J, Kaadige MR, Rao A, Sheppard KE, Hugo W, Pupo GM, et al (2014) Response of BRAF-mutant melanoma to BRAF inhibition is mediated by a network of transcriptional regulators of glycolysis. Cancer Discov 4: 423–433

Patra KC, Wang Q, Bhaskar PT, Miller L, Wang Z, Wheaton W, Chandel N, Laakso M, Muller WJ, Allen EL, et al (2013) Hexokinase 2 is required for tumor initiation and maintenance and its systemic deletion is therapeutic in mouse models of cancer. Cancer Cell 24: 213–228

Qin J-Z, Xin H & Nickoloff BJ (2010) Targeting glutamine metabolism sensitizes melanoma cells to TRAIL-induced death. Biochemical and Biophysical Research Communications 398: 146–152

Queiroz RML, Smith T, Villanueva E, Marti-Solano M, Monti M, Pizzinga M, Mirea D-M, Ramakrishna M, Harvey RF, Dezi V, et al (2019) Comprehensive identification of RNA–protein interactions in any organism using orthogonal organic phase separation (OOPS). Nat Biotechnol 37: 169–178

Rosenbaum SR, Caksa S, Stefanski CD, Trachtenberg IV, Wilson HP, Wilski NA, Ott CA, Purwin TJ, Haj JI, Pomante D, et al (2024) SOX10 Loss Sensitizes Melanoma Cells to Cytokine-Mediated Inflammatory Cell Death. Molecular Cancer Research 22: 209–220

Rossman MG, Liljas A, Brändén C-I & Banaszak LJ (1975) 2 Evolutionary and Structural Relationships among Dehydrogenases. In The Enzymes, Boyer PD (ed) pp 61–102. Academic Press

Scott DA, Richardson AD, Filipp FV, Knutzen CA, Chiang GG, Ronai ZA, Osterman AL & Smith JW (2011) Comparative metabolic flux profiling of melanoma cell lines: beyond the Warburg effect. J Biol Chem 286: 42626–42634

Servant N, Philippe LR, phupe & Allain F (2023) bioinfo-pf-curie/RNA-seq: v4.0.0.

Shan W, Zhou Y & Tam KY (2022) The development of small-molecule inhibitors targeting hexokinase 2. Drug Discovery Today 27: 2574–2585

Shen S, Faouzi S, Bastide A, Martineau S, Malka-Mahieu H, Fu Y, Sun X, Mateus C, Routier E, Roy S, et al (2019) An epitranscriptomic mechanism underlies selective mRNA translation remodelling in melanoma persister cells. Nat Commun 10: 5713

Simsek D, Tiu GC, Flynn RA, Byeon GW, Leppek K, Xu AF, Chang HY & Barna M (2017) The Mammalian Ribo-interactome Reveals Ribosome Functional Diversity and Heterogeneity. Cell 169: 1051–1065.e18

Song P, Yang F, Jin H & Wang X (2021) The regulation of protein translation and its implications for cancer. Sig Transduct Target Ther 6: 1–9

Trendel J, Schwarzl T, Horos R, Prakash A, Bateman A, Hentze MW & Krijgsveld J (2019) The Human RNA-Binding Proteome and Its Dynamics during Translational Arrest. Cell 176: 391–403.e19

Urdaneta EC, Vieira-Vieira CH, Hick T, Wessels H-H, Figini D, Moschall R, Medenbach J, Ohler U, Granneman S, Selbach M, et al (2019) Purification of cross-linked RNA-protein complexes by phenol-toluol extraction. Nat Commun 10: 990

Vander Heiden MG, Cantley LC & Thompson CB (2009) Understanding the Warburg Effect: The Metabolic Requirements of Cell Proliferation. Science 324: 1029–1033

Wang F, Zhang S, Vuckovic I, Jeon R, Lerman A, Folmes CD, Dzeja PP & Herrmann J (2018) Glycolytic Stimulation Is Not a Requirement for M2 Macrophage Differentiation. Cell Metabolism 28: 463–475.e4

Wang W, Liu Z, Zhao L, Sun J, He Q, Yan W, Lu Z & Wang A (2017) Hexokinase 2 enhances the metastatic potential of tongue squamous cell carcinoma via the SOD2-H2O2 pathway. Oncotarget 8: 3344–3354

Warburg O (1956) On Respiratory Impairment in Cancer Cells. Science 124: 269–270

Weinberg F, Hamanaka R, Wheaton WW, Weinberg S, Joseph J, Lopez M, Kalyanaraman B, Mutlu GM, Budinger GRS & Chandel NS (2010) Mitochondrial metabolism and ROS generation are essential for Kras-mediated tumorigenicity. Proceedings of the National Academy of Sciences 107: 8788–8793

Wickham H (2009) ggplot2: Elegant Graphics for Data Analysis New York, NY: Springer

Wolf A, Agnihotri S, Micallef J, Mukherjee J, Sabha N, Cairns R, Hawkins C & Guha A (2011) Hexokinase 2 is a key mediator of aerobic glycolysis and promotes tumor growth in human glioblastoma multiforme. J Exp Med 208: 313–326

Wolf AJ, Reyes CN, Liang W, Becker C, Shimada K, Wheeler ML, Cho HC, Popescu NI, Coggeshall KM, Arditi M, et al (2016) Hexokinase Is an Innate Immune Receptor for the Detection of Bacterial Peptidoglycan. Cell 166: 624–636

Xiao Z, Zou Q, Liu Y & Yang X (2016) Genome-wide assessment of differential translations with ribosome profiling data. Nat Commun 7

Zhang C, Liu J, Liang Y, Wu R, Zhao Y, Hong X, Lin M, Yu H, Liu L, Levine AJ, et al (2013) Tumour-associated mutant p53 drives the Warburg effect. Nat Commun 4: 2935

Zhang W, Xie M, Shu M-D, Steitz JA & DiMaio D (2016) A proximity-dependent assay for specific RNA–protein interactions in intact cells. RNA 22: 1785–1792

Zheng Y, Zhan Y, Zhang Y, Zhang Y, Liu Y, Xie Y, Sun Y, Qian J, Ding Y, Ding Y, et al (2023) Hexokinase 2 confers radio-resistance in hepatocellular carcinoma by promoting autophagy-dependent degradation of AIMP2. Cell Death Dis 14: 1–13

Zhou Y, Yi X, Stoffer JB, Bonafe N, Gilmore-Hebert M, McAlpine J & Chambers SK (2008) The Multifunctional Protein Glyceraldehyde-3-Phosphate Dehydrogenase Is Both Regulated and Controls Colony-Stimulating Factor-1 Messenger RNA Stability in Ovarian Cancer. Molecular Cancer Research 6: 1375–1384

