## Supplementary Figures 1-7 for "An extra-glycolytic function for hexokinase 2 as an RNA-binding protein regulating *SOX10* mRNA translation in melanoma"

**SUPPLEMENTARY INFORMATION**

**Figures S1-S7**

### An extra-glycolytic function for hexokinase 2 as an RNA-binding protein regulating *SOX10* mRNA translation in melanoma

**Ana Luisa Dian, Antoine Moya-Plana, Giuseppina Claps, Céline M. Labbé, Virginie Quidville, Dorothée Baille, Virginie Raynal, Sylvain Baulande, Caroline Robert, Stéphan Vagner**

### Figure S1

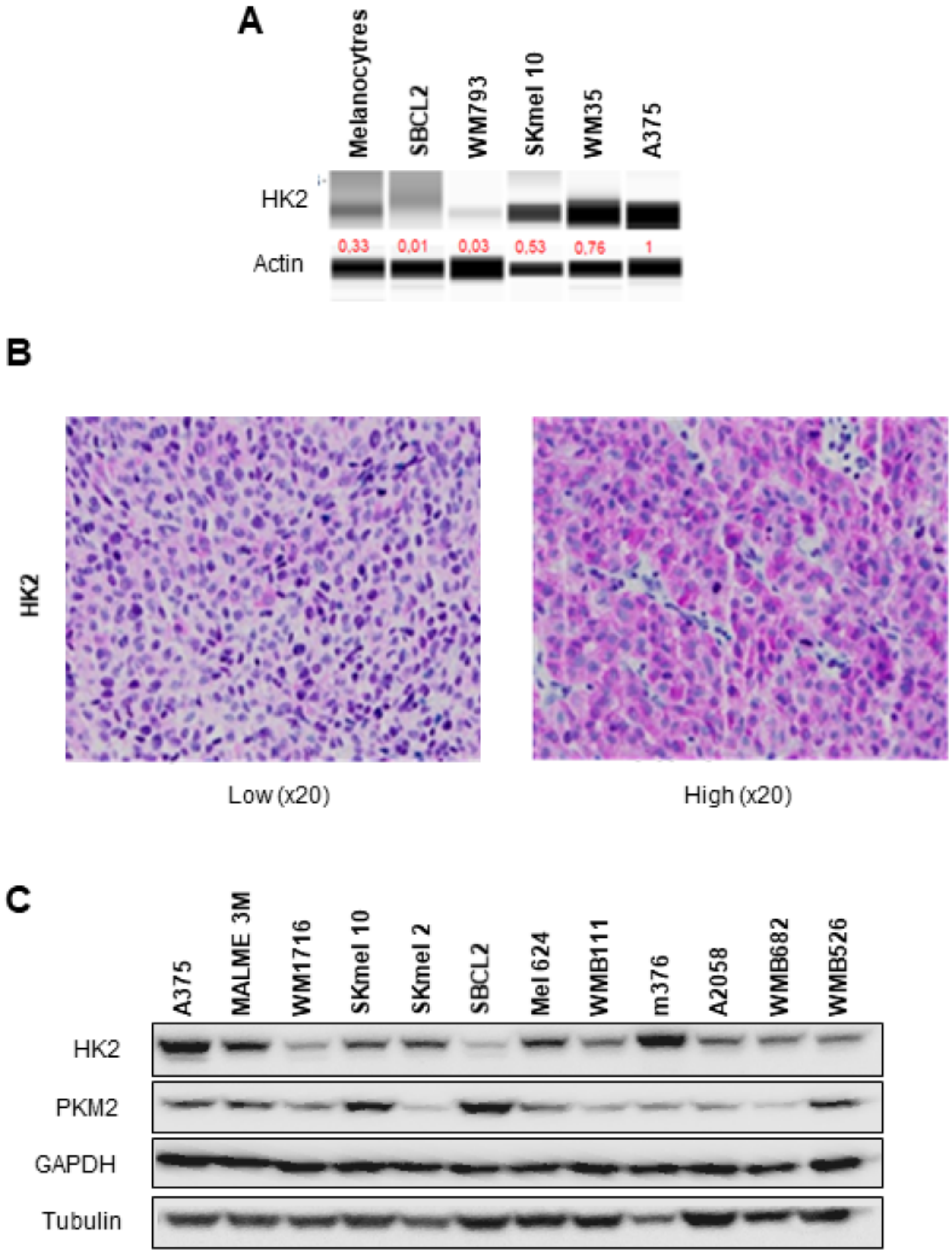
A
Melanocytres
SBCL2
WM793
SKmel 10
WM35
A375
HK2
Actin
0,33
0,01
0,03
0,53
0,76
1
B
HK2
Low (x20)
High (x20)
C
A375
MALME 3M
WM1716
SKmel 10
SKmel 2
SBCL2
Mel 624
WMB111
m376
A2058
WMB682
WMB526
HK2
PKM2
GAPDH
Tubulin

**Figure S1. HK2 expression in melanoma cell lines and tumours.**

**(A)** HK2 expression level assessed by automated capillary western blot (Protein Simple ®) analysis of normal melanocytes, early superficial melanoma with radial growth (RGP, SBCL2), early invasive melanoma (VGP, WM793), low invasive (SKMel10) and high invasive (A375) metastatic melanoma cell lines. Actin was used as loading control. **(B)** Immunohistochemical analysis of HK2 expression in two representative samples from a cohort of 31 patients with cutaneous melanoma. **(C)** Western blot analysis of key glycolytic enzymes (HK2, PKM2 and GAPDH) in a variety of melanoma cell lines. Tubulin was used as loading control.

#### Figure S2

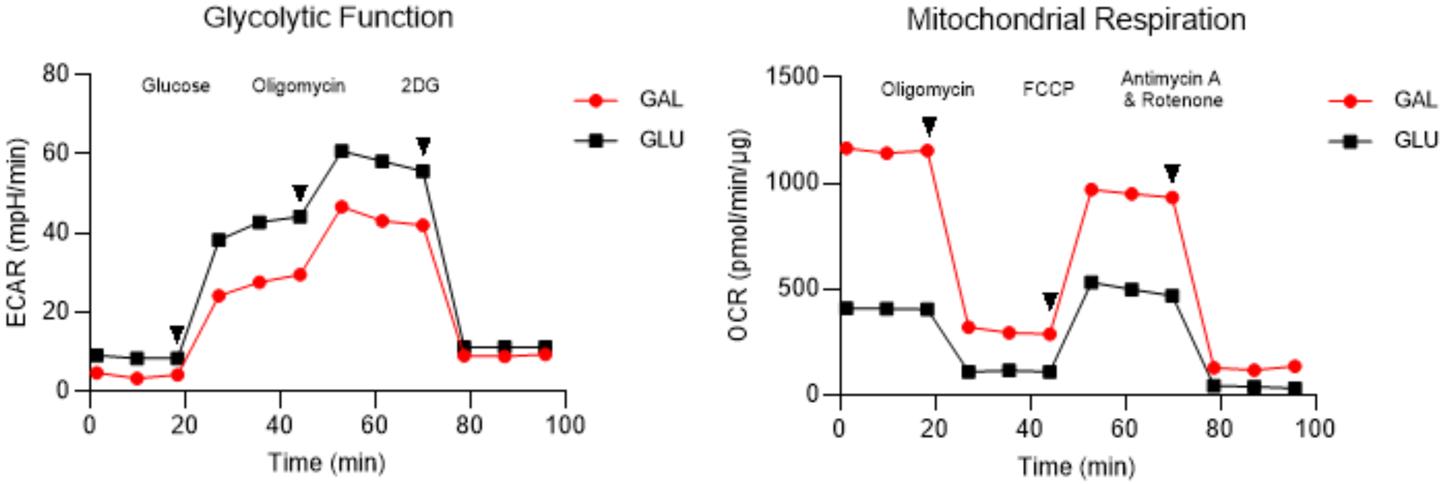


**Metabolic profile of A375 cells analysed on a XF96 Extracellular Flux Analyzer.**

Left panel: the extra-cellular acidification rate (ECAR), an indicator of aerobic glycolysis, was measured in A375 cells grown in either glucose or galactose-containing medium followed by consecutive treatments of Glucose, oligomycin, and 2-DG. Right panel: the oxygen consumption rate (OCR) was measured in A375 cells cultured either in glucose or galactose-containing medium followed by consecutive treatments of oligomycin, FCCP, and antimycin A & rotenone.

#### Figure S3

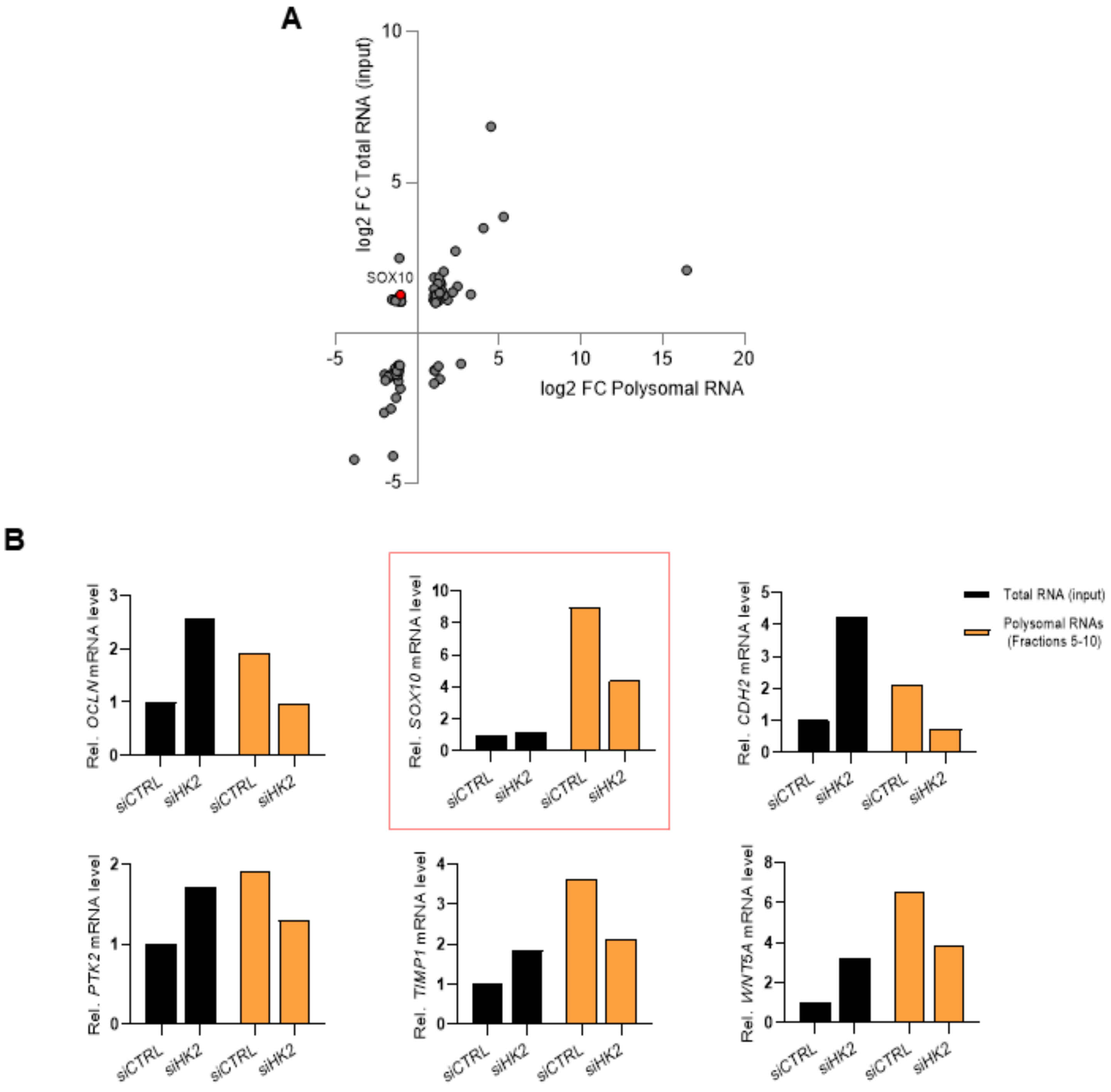
A
log2 FC Total RNA (input)
log2 FC Polysomal RNA
SOX10
B
Rel. OCLN mRNA level
Rel. SOX10 mRNA level
Rel. CDH2 mRNA level
Rel. PTK2 mRNA level
Rel. TIMP1 mRNA level
Rel. WNT5A mRNA level
siCTRL
siHK2
Total RNA (input)
Polysomal RNAs
(Fractions 5-10)

**Figure S3. HK2 regulates the expression of cancer-related genes.**

**(A)** RT-qPCR quantification of 84 EMT-associates genes. Transcriptional (total RNA) and translational (polysomal RNA) levels of each gene were obtained from polysome profiling of A375 cells transfected with siRNAs targeting HK2 (siHK2) or control (siCTR) and measured using $RT^2$ Profiler PCR Array. **(B)** RT-qPCR analysis of potential key genes of EMT (*OCLN*, *SOX10*, *CDH2*, *PTK2*, *TIMP1*, and *WNT5A*) whose expression could be translationally regulated by HK2. Transcriptional (total RNA; black) and translational (polysomal RNA; orange) levels of each gene were obtained from polysome profiling of A375 cells transfected with siRNAs targeting HK2 (siHK2) or control (siCTR) and measured using $RT^2$ Profiler PCR Array.

#### Figure S4

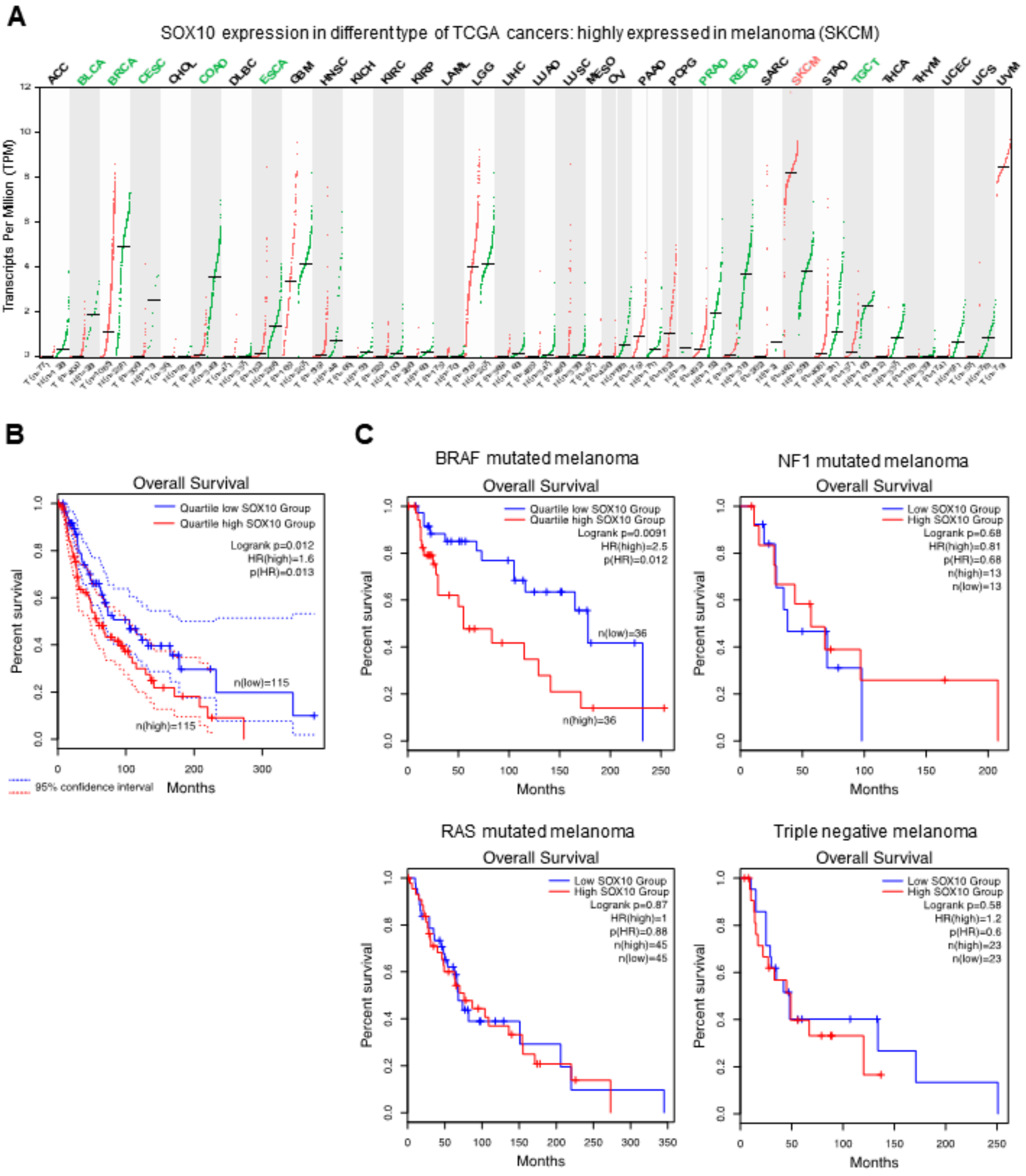
A
SOX10 expression in different type of TCGA cancers: highly expressed in melanoma (SKCM)
ACC
BLCA
BRCA
CESC
CHOL
COAD
DLBC
ESCA
GBM
HNSC
KICH
KIRC
KIRP
LAML
LGG
LIHC
LUAD
LUSC
MESO
OV
PAAD
PCPG
PRAD
READ
SARC
SKCM
STAD
TGCT
THCA
THYM
UCEC
UCS
UVM
Transcripts Per Million (TPM)
B
Overall Survival
Quartile low SOX10 Group
Quartile high SOX10 Group
Logrank p=0.012
HR(high)=1.6
p(HR)=0.013
n(low)=115
n(high)=115
Percent survival
Months
95% confidence interval
C
BRAF mutated melanoma
Overall Survival
Quartile low SOX10 Group
Quartile high SOX10 Group
Logrank p=0.0091
HR(high)=2.5
p(HR)=0.012
n(low)=36
n(high)=36
Percent survival
Months
NF1 mutated melanoma
Overall Survival
Low SOX10 Group
High SOX10 Group
Logrank p=0.68
HR(high)=0.81
p(HR)=0.68
n(high)=13
n(low)=13
Percent survival
Months
RAS mutated melanoma
Overall Survival
Low SOX10 Group
High SOX10 Group
Logrank p=0.87
HR(high)=1
p(HR)=0.88
n(high)=45
n(low)=45
Percent survival
Months
Triple negative melanoma
Overall Survival
Low SOX10 Group
High SOX10 Group
Logrank p=0.58
HR(high)=1.2
p(HR)=0.6
n(high)=23
n(low)=23
Percent survival
Months

**Figure S4. SOX10 expression is associated with prognosis in patients with cutaneous melanoma.**

**(A)** SOX10 expression in different type of solid cancers from TGCA database. **(B)** Analysis of the impact of SOX10 expression in overall survival of patients with cutaneous melanoma. **(C)** Analysis of the impact of SOX10 expression in overall survival according to molecular profile of the tumour. T: tumour (red dots); N: normal tissue (green dots).

### Figure S5

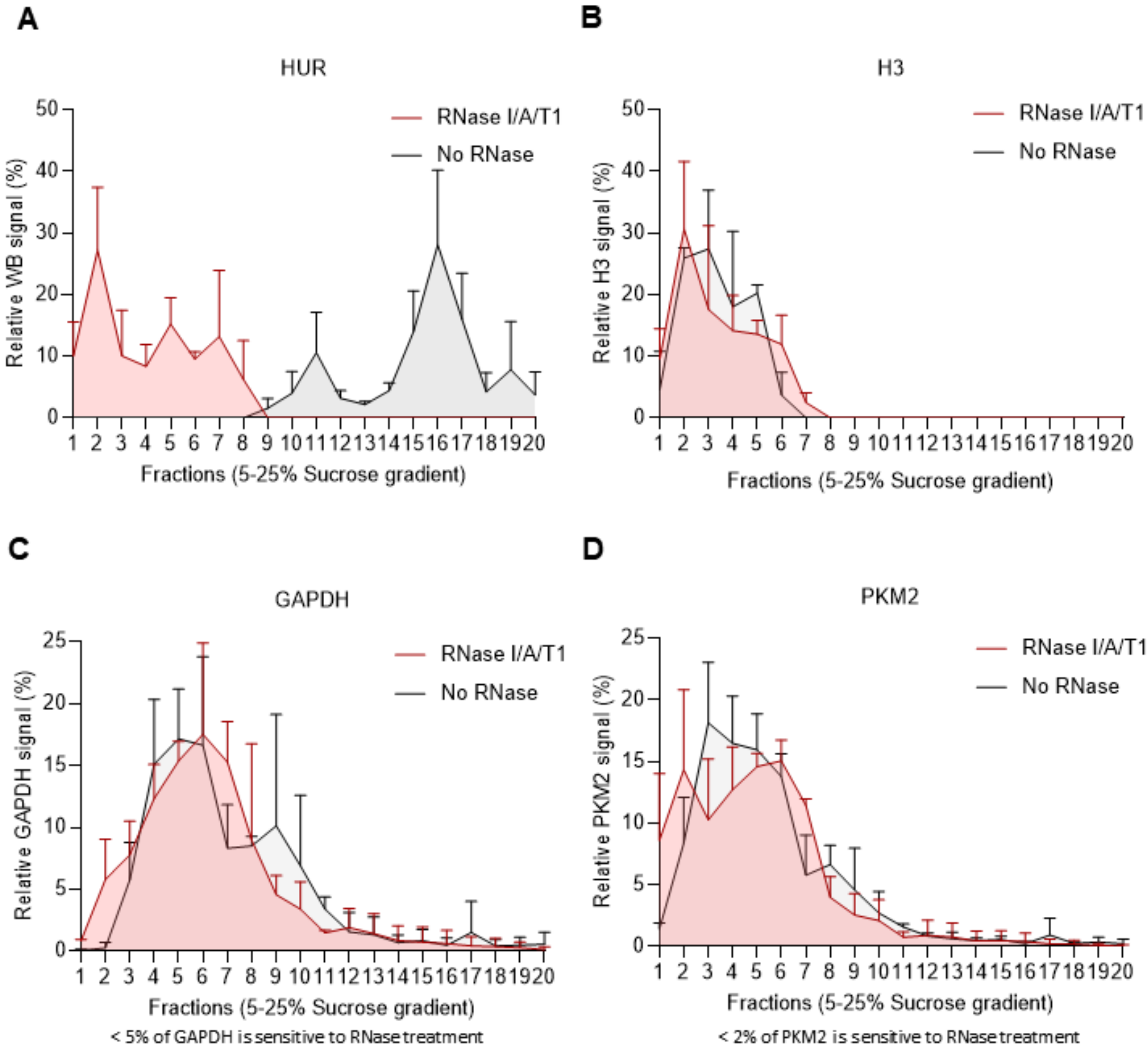


**Sucrose density gradient centrifugation and fractionation of A375 cell lysates.**

Western blot quantification representing the % of **(A)** HUR, **(B)** H3, **(C)** GAPDH and **(D)** PKM2 in each sucrose fraction of lysates treated with RNase I/A/T1 or left untreated (SD, n = 3 biological replicates).

#### Figure S6

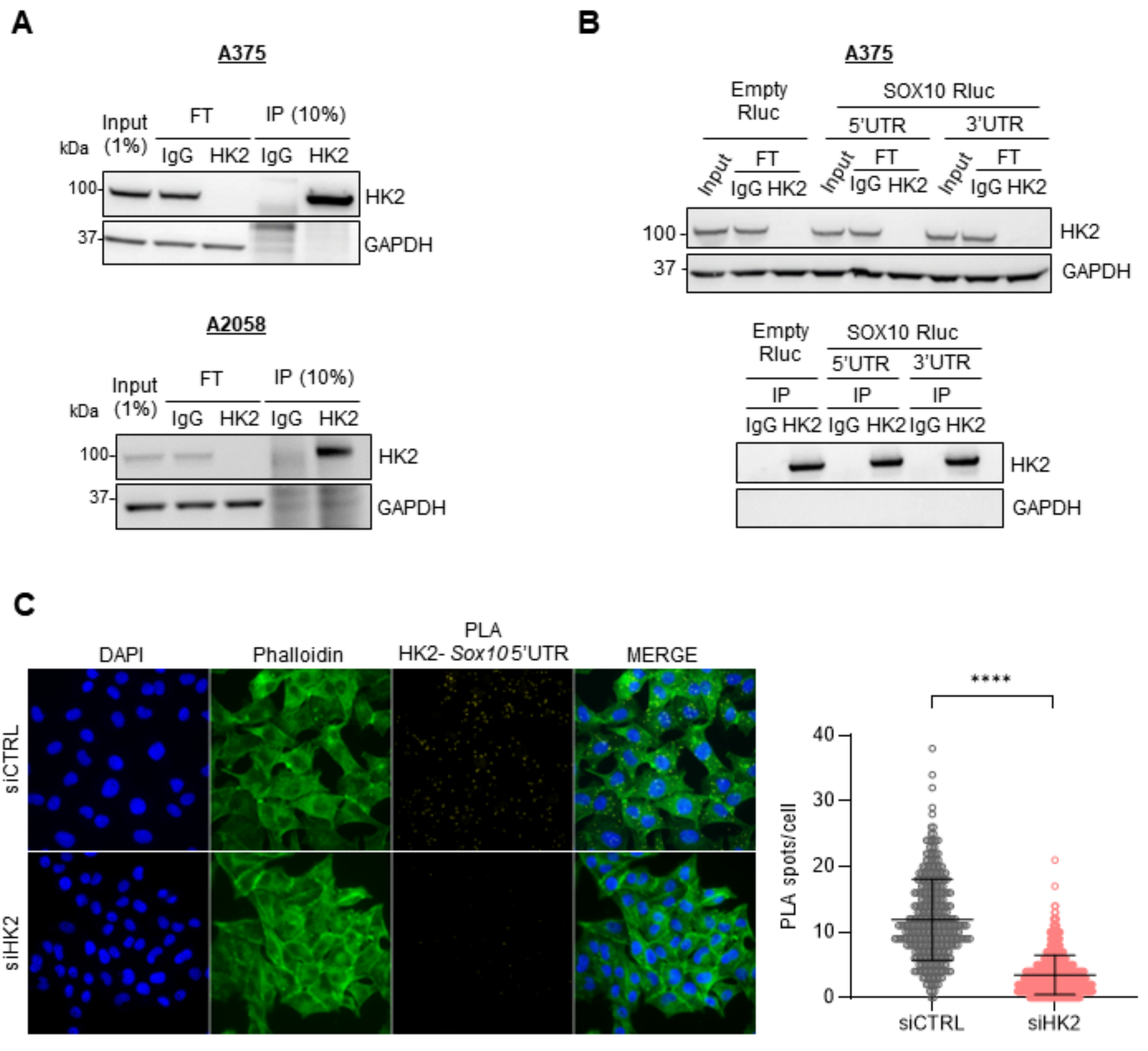
A
A375
Input (1%)
FT
IP (10%)
kDa
IgG
HK2
IgG
HK2
100
37
HK2
GAPDH
A2058
Input (1%)
FT
IP (10%)
kDa
IgG
HK2
IgG
HK2
100
37
HK2
GAPDH
B
A375
Empty Rluc
SOX10 Rluc
5'UTR
3'UTR
Input
FT
IgG HK2
100
37
HK2
GAPDH
Empty Rluc
SOX10 Rluc
5'UTR
3'UTR
IP
IgG HK2
HK2
GAPDH
C
DAPI
Phalloidin
PLA
HK2- Sox10 5'UTR
MERGE
siCTRL
siHK2
****
PLA spots/cell
40
30
20
10
0
siCTRL
siHK2

**Figure S6. Analysis of HK2-*SOX10* mRNA interaction.**

**(A)** Western blot of HK2 and GAPDH (normalization control) from RIP experiment performed on A375 and A2058 melanoma cell lines (representative images, n = 3 biological replicates). **(B)** Western blot of HK2 and GAPDH (normalization control) from RIP experiment performed on A375 cells transfected with luciferase reporters containing the *SOX10* 5'UTR and 3'UTR sequences upstream of the Renilla luciferase reporter gene. An empty reporter was used as control (representative images, n = 3 biological replicates). **(C)** Representative fluorescence images and quantification of RNA-proximity ligation assays (PLA) of HK2 protein and *SOX10* mRNA in A375 cells transfected with siRNAs targeting HK2 (siHK2, light red) or control (siCTRL, grey) (n = 3 biological replicates). After 48h, cells were fixed, permeabilized, and incubated with anti-sense *SOX10* 5'UTR oligonucleotide probes and anti-HK2 antibody. Scale bar, 10 µm. HK2-*SOX10* mRNA PLA signal (yellow), Phalloidin staining (green), and DAPI staining (blue). The significance for PLA values was derived from the Mann-Whitney statistical test (SD, **** $p \leq 0.0001$).

#### Figure S7

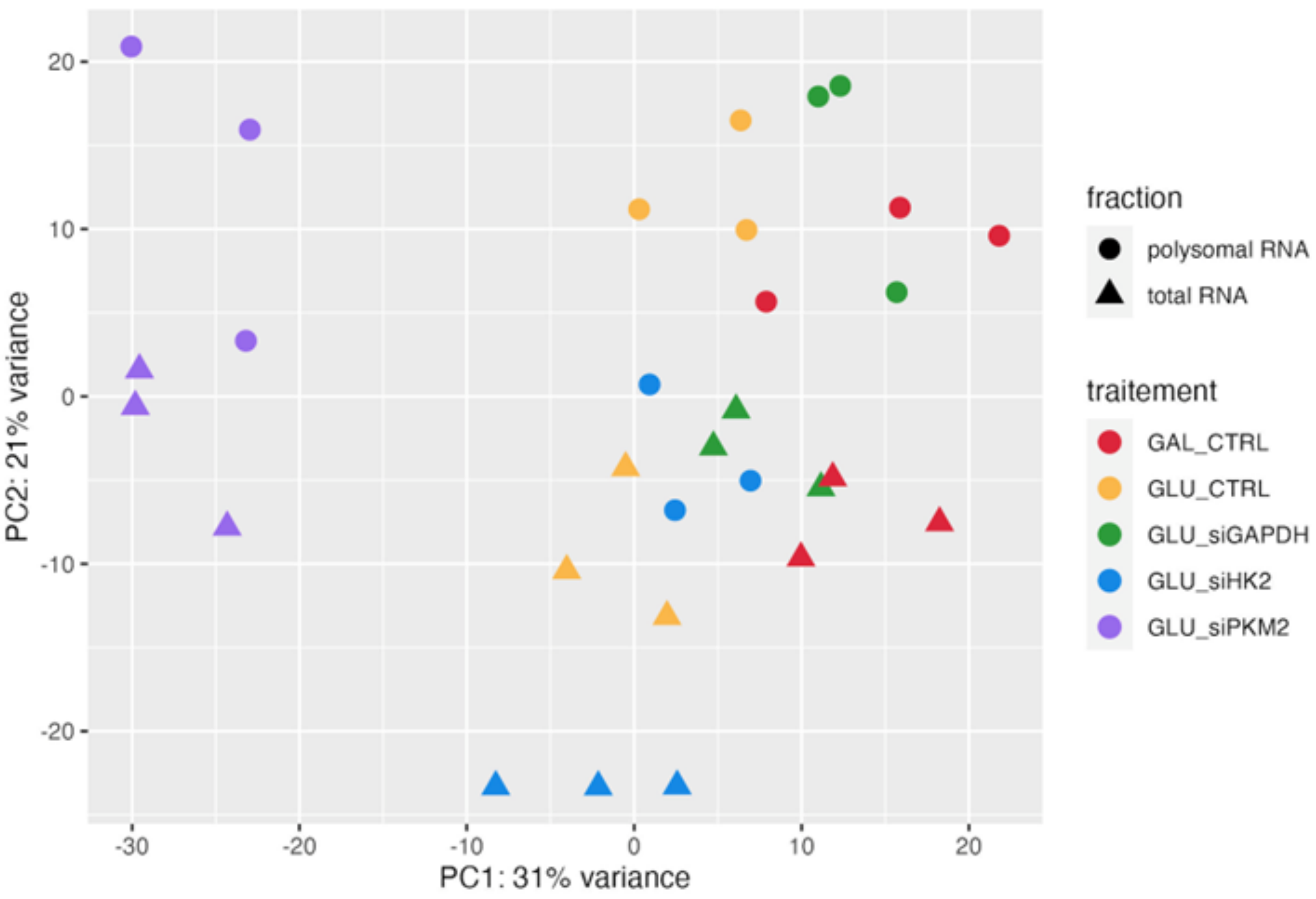


**PCA plot using the VST data.**

The variance stabilizing transformation (VST) offered by DESeq2 was used on the raw count data to stabilize the variance across the mean. The principal components analysis (PCA) plot was built using the ggplot2 package.
