## Supplementary Tables S2-S3 for "An extra-glycolytic function for hexokinase 2 as an RNA-binding protein regulating *SOX10* mRNA translation in melanoma"

**SUPPLEMENTARY INFORMATION**

**Tables S2 and S3**

### An extra-glycolytic function for hexokinase 2 as an RNA-binding protein regulating *SOX10* mRNA translation in melanoma

**Ana Luisa Dian, Antoine Moya-Plana, Giuseppina Claps, Céline M. Labbé, Virginie Quidville, Dorothée Baille, Séverine Roy, Virginie Raynal, Sylvain Baulande, Caroline Robert, Stéphan Vagner**

**Table S2. Transcriptome and translatoma analyses on RT$^2$ Profiler$^{tm}$ PCR Array.** HP, heavy polysomes.

| Gene Name | **Input siHK2 vs Input siCTRL** log2(fold change) | **HP siHK2 vs HP siCTRL** log2(fold change) |
|---|---|---|
| AHNAK | 1.06 | -1.35 |
| AKT1 | -1.08 | -1.09 |
| BMP1 | 1.33 | 1.34 |
| BMP2 | 2.04 | 1.59 |
| BMP7 | -1.68 | 1 |
| CALD1 | 1 | 1.1 |
| CAMK2N1 | -1.27 | -1.28 |
| CAV2 | -1.28 | -1.16 |
| CDH1 | 1.05 | -1.01 |
| CDH2 | 1.09 | -1.38 |
| COL1A2 | 2.48 | -1.12 |
| COL3A1 | 2.08 | 16.46 |
| COL5A2 | 1.24 | 1.13 |
| CTNNB1 | -1.38 | -1.33 |
| DESI1 | 1.42 | 1.47 |
| DSC2 | -1.41 | -1.2 |
| DSP | 1.08 | 1.02 |
| EGFR | -1.58 | -1.97 |
| ERBB3 | 1.24 | 1 |
| ESR1 | -2.16 | -1.32 |
| F11R | -1.46 | -1.81 |
| FGFBP1 | 1.46 | 1.01 |
| FN1 | 1.64 | 1.25 |
| FOXC2 | 3.86 | 5.25 |
| FZD7 | 1.49 | 1.14 |
| GEMIN2 | -1.61 | -1.19 |
| GNG11 | 1.15 | -1.2 |
| GSC | 1.21 | -1.07 |
| GSK3B | 1.15 | -1.08 |
| IGFBP4 | -1.11 | -1.14 |
| IL1RN | 1.09 | 1.01 |
| ILK | -1.4 | -1.52 |
| ITGA5 | 1.3 | 1.11 |
| ITGAV | 1.08 | 1.17 |
| ITGB1 | 1.37 | 1.3 |
| JAG1 | 1.36 | 1.14 |
| KRT14 | -2.52 | -1.63 |
| KRT19 | 1.09 | 1.05 |
| KRT7 | 1.09 | 1.01 |

| | | |
|---|---|---|
| MAP1B | 1.84 | 1.31 |
| MMP2 | 1.31 | 1.36 |
| MMP3 | -1.54 | 1.36 |
| MMP9 | 1.03 | -1.05 |
| MSN | 1.69 | 1.39 |
| MST1R | -1.03 | 2.66 |
| NODAL | 1.35 | 2.14 |
| NOTCH1 | 1.45 | 1.03 |
| NUDT13 | 1.04 | -1.02 |
| OCLN | 1.11 | -1.57 |
| PDGFRB | 2.72 | 2.31 |
| PLEK2 | 1.28 | 3.26 |
| PTK2 | 1.15 | -1.17 |
| PTP4A1 | -1.08 | -1.18 |
| RAC1 | 1.09 | 1.33 |
| RGS2 | 3.48 | 4.03 |
| SERPINE1 | 6.86 | 4.49 |
| SMAD2 | 1.06 | 1.23 |
| SNAI1 | 1.61 | 1.44 |
| SNAI2 | -2.66 | -2.05 |
| SNAI3 | 1.54 | 2.43 |
| SOX10 | 1.27 | -1.04 |
| SPARC | 1.15 | 1.1 |
| SPP1 | 1.24 | 1.64 |
| STAT3 | -1.48 | -1.73 |
| STEAP1 | -1.11 | 1.26 |
| TCF3 | 1.05 | -1.12 |
| TCF4 | -1.85 | -1.06 |
| TFP12 | 1.09 | 1.83 |
| TGFB1 | 1.44 | 1.41 |
| TGFB2 | 1.84 | 1.01 |
| TGFB3 | 1.02 | 1.09 |
| TIMP1 | 1.2 | -1.03 |
| TMEFF1 | -1.26 | 1.06 |
| TMEM132A | 1.11 | 1.05 |
| TSPAN13 | -1.25 | 1.02 |
| TWIST1 | -1.34 | -1.3 |
| VCAN | -4.22 | -3.89 |
| VIM | 1.04 | 1.17 |
| VPS13A | -1.38 | -1.67 |
| WNT11 | -1.38 | -2.01 |
| WNT5A | 1.04 | -1.03 |
| WNT5B | -4.1 | -1.51 |
| ZEB1 | -1.18 | -1.38 |

| | | |
|---|---|---|
| ZEB2 | -1.25 | -1.17 |

**Table S3. Relevant oligonucleotides.**

| NAME OF THE OLIGONUCLEOTIDE | SEQUENCE |
|---|---|
| **RTqPCR primers** | |
| VCL Forward | TGATGATTAGAGACATCACCGCT |
| VCL Reverse | AACAAAGGAGAAACCTGACCT |
| GAPDH Forward | TCCCATCACCATCTTCCAGG |
| GAPDH Reverse | TCCATGGTGGTGAAGACGC |
| TBP Forward | GGAAGGGGCATTATTTGTG |
| TBP Reverse | GCCCAGATAGCAGCACGGTA |
| ACT Forward | CCGTGTTTCCTTCCATCGTC |
| ACT Reverse | ACGATGCCATGCTCAATGGG |
| Renilla Forward | ACGGATGATAACTGGTCCGC |
| Renilla Reverse | CGCGCTACTGGCTCAATATG |
| Firefly Forward | GAAATGTCCGTTCGGTTGGC |
| Firefly Reverse | TCCGATAAATAACGCGCCCA |
| OCLN Forward | GGTCGGGCCCAGTTGC |
| OCLN Reverse | ATGATTCGGTTTGAATTCATCAGG |
| CDH2 Forward | ACTCCAGGGGACCTTTTCCT |
| CDH2 Reverse | TGCCCTCAAATGAAACCGGG |
| PTK2 Forward | TTGGGCGGAAAGAAATCCTG |
| PTK2 Reverse | GTCCAGGTTGGCAGTAGGAG |
| TMP1 Forward | GACACCAGAGAACCCACCAT |
| TMP1 Reverse | CACGAACTTGGCCCTGATGA |
| WNT5A Forward | GCTCGCATCCTCATGAACCT |
| WNT5A Reverse | GCCACATCAGCCAGGTTGTA |
| SOX10 5'UTR Forward | CACTTCCTAAGGACGAGCCC |
| SOX10 5'UTR Reverse | TCCTCGCAAAGAGTCCAACG |
| SOX10 CDS Forward | GGCTGCTGAACGAAAGTGA |
| SOX10 CDS Reverse | TCTTGTAGTGGGCCTGGATG |
| SOX10 POLYSOME Forward | CCAGGCCCACTACAAGAGC |
| SOX10 POLYSOME Reverse | CTCTGTCTTCGGGGTGGTTG |
